# Hybrid dedicated and distributed coding in PMd/M1 provides separation and interaction of bilateral arm signals

**DOI:** 10.1101/2020.09.23.310664

**Authors:** Tanner C. Dixon, Christina M. Merrick, Joni D. Wallis, Richard B. Ivry, Jose M. Carmena

## Abstract

Pronounced activity is observed in both hemispheres of the motor cortex during preparation and execution of unimanual movements. The organizational principles of bi-hemispheric signals and the functions they serve throughout motor planning remain unclear. Using an instructed-delay reaching task in monkeys, we identified two components in population responses spanning PMd and M1. A “dedicated” component, which segregated activity at the level of individual units, emerged in PMd during preparation. It was most prominent following movement when M1 became strongly engaged, and principally involved the contralateral hemisphere. In contrast to recent reports, these dedicated signals solely accounted for divergence of arm-specific neural subspaces. The other “distributed” component mixed signals for each arm within units, and the subspace containing it did not discriminate between arms at any stage. The statistics of the population response suggest two functional layers of the cortical network: one that spans both hemispheres for supporting preparatory and ongoing processes, and another that is predominantly housed in the contralateral hemisphere and specifies unilateral output.

## INTRODUCTION

In the primate cortex, direct control of arm movement is primarily mediated by contralateral descending projections (Lawrence and Kuypers, 1968; Brinkman and Kuypers, 1973; Soteropoulos et al., 2011). However, numerous studies have observed activity changes in the motor cortex during movements of the ipsilateral arm (Matsunami and Hamada, 1981; Donchin et al., 1998; Hoshi and Tanji, 2002; Steinberg et al., 2002; Cisek et al., 2003; Ganguly et al., 2009; Ames and Churchland, 2019; Heming et al., 2019) and hand (Tanji et al., 1988; Verstynen et al., 2005; Diedrichsen et al., 2013). The functional role of this ipsilateral activity has been the subject of considerable debate, with hypotheses ranging from a role in postural support, bimanual coordination, or an extrapyramidal control signal for unimanual movements.

Neurons in the primate dorsal premotor cortex (PMd) play a critical role in motor preparation (Weinrich et al., 1984; Shen and Alexander, 1997; Tanji et al., 1988; Hoshi and Tanji, 2002; Cisek et al., 2003). Interestingly, their response properties and degree of laterality appear to change across the course of preparation. For example, within PMd, individual units exhibit a transition from effector-independent to effector-dependent encoding between preparatory and execution phases of reaching. In contrast, units in primary motor cortex (M1) mainly become active during movement itself and show a pronounced contralateral bias (Cisek et al., 2003). This suggests a transition from abstract planning to explicit specification of motor output parameters in the signals of individual neurons. A similar transition has been shown in the activation of different cell-types within rodent motor areas (Li et al., 2015; Soma et al., 2017). These studies have found that neurons with intracortical projections show little lateral bias, particularly during pre-movement phases. In contrast, neurons with descending output display much stronger laterality, especially just before and during movement. This adds yet another level of granularity in the discussion of lateralized motor function. Collectively, these single-unit studies support a notion that there exist two distinct components within the motor cortex: one that is bilateral and likely involved in abstract processing, and another that is dedicated to a single side of the body for execution.

The classical perspectives outlined above have been revisited in studies that focus on population-level analysis, considering instead how computations might be reflected in the way the network coordinates activity. Low-dimensional representations of large-scale neural recordings can be used to characterize these network patterns, revealing changes in covariance structure across behavioral settings that are not evident when looking at single neurons in isolation (Cunningham and Yu, 2014). Ostensibly, these changes reflect reorganization of the population as it engages in different computational processes. Using these methods, pre-movement activity has been shown to evolve within an “output-null” subspace towards an optimal initial population state (Churchland et al., 2010; Kaufman et al., 2014; Elsayed et al., 2016). This initial state is advantageously positioned for engaging the internal dynamics of the network to produce patterned activity in an “output-potent” subspace for driving movement (Churchland et al., 2012; Shenoy et al., 2013; Sussillo et al., 2015). There is some evidence that bilateral activity may support these preparatory and dynamic properties (Li et al., 2016). Similar to the output-null and output-potent subspaces, arm-specific subspaces have also been observed in M1 during rhythmic movements (Ames and Churchland, 2019) and in response to joint perturbations (Heming et al., 2019). It remains unclear precisely what organizational principles produce these arm-specific subspaces, whether signals are fully separated at the level of the population, and how such properties develop across preparation and movement.

There are two fundamental and mutually non-exclusive ways that population signals may specify the selected arm across preparation and movement. (1) Signals may consolidate within dedicated sub-populations for each arm (i.e., within hemispheres, brain areas, or cell-types). (2) Signals for each arm may be distributed across the same units yet maintain unique covariance structure that separates them along arm-specific neural dimensions. Importantly, either of these architectures provides a way for downstream targets to discriminate signals and also yields the mathematical result of divergent subspaces. The first method is necessarily true: contralateral biases have been consistently observed during movement and, to a lesser extent, during preparation as well. Such lateral biases will trivially orthogonalize arm signals. The question is whether the second method is also true. Either signals that are mixed within units become separated (arm-specific) in the population readout, or they exist within a space where the same patterns of activity are involved in computations relevant to both arms. This is a vital distinction to make, as it constrains the possible roles that bilateral activity can play at each stage of processing and may point to an important heterogeneity in the population statistics.

In the present study, we recorded large populations of single-units in PMd and M1 bilaterally while monkeys performed an instructed-delay unimanual reaching task. As activity emerged during preparation, there was a tendency for units with stronger arm preference to be more highly modulated, therefore representing a larger proportion of the population variance. As a result, the signals for each arm were largely segregated, primarily within contralateral PMd. During the transition to movement, M1 became more prominently involved and the signals for each arm became increasingly segregated. This unit-level segregation caused the subspaces corresponding to each arm to diverge across the trial. However, we did observe target-specific information that was mixed within single-units, indicating incomplete separation of arm signals. Importantly, the subspace containing this information did not segregate signals for the two arms at any stage. Taken together, the results point to two primary components in the population response: (1) A dedicated component that develops across preparation, reaches a maximum during movement, and mirrors the lateralized anatomy of corticospinal output with its contralateral bias. (2) A distributed component that represents far less variance, particularly during movement, and provides a space in which bilateral arm signals may easily interact.

## RESULTS

### Behavior

Two macaque monkeys were trained to perform an instructed-delay reaching task in 3-D space (Figure 1A). Reaching movements were freely performed in an open area while kinematics were recorded using optical motion tracking. Visual feedback of endpoint position and task cues were provided through a virtual 3-D display. Each trial had three phases (Figure 1B). For the Rest phase, the monkey placed both hands in start targets positioned near the torso and remained still for 500 ms. For the Instruct phase, an instructional cue appeared at one of six target locations. The color of the cue specified the required hand for the forthcoming trial. The monkey was required to keep both hands in the rest positions while the cue remained visible for a variable interval (500 – 1500 ms). The Move phase was initiated when the start position marker for the reaching hand disappeared and the cue at the target location increased in size, which signaled the animal to reach. The monkey received a juice reward if it accurately reached the target and maintained the final position for 250 ms, while keeping the non-cued hand at its start position for the duration of the trial. 300ms representative windows from each phase were used in data analysis. Trials were blocked for each arm, with each block consisting of 2 trials per target in a randomized order (i.e. alternating 2 trials per target for the left arm, then 2 trials per target for the right; Figure 1C).

**Figure 1.**
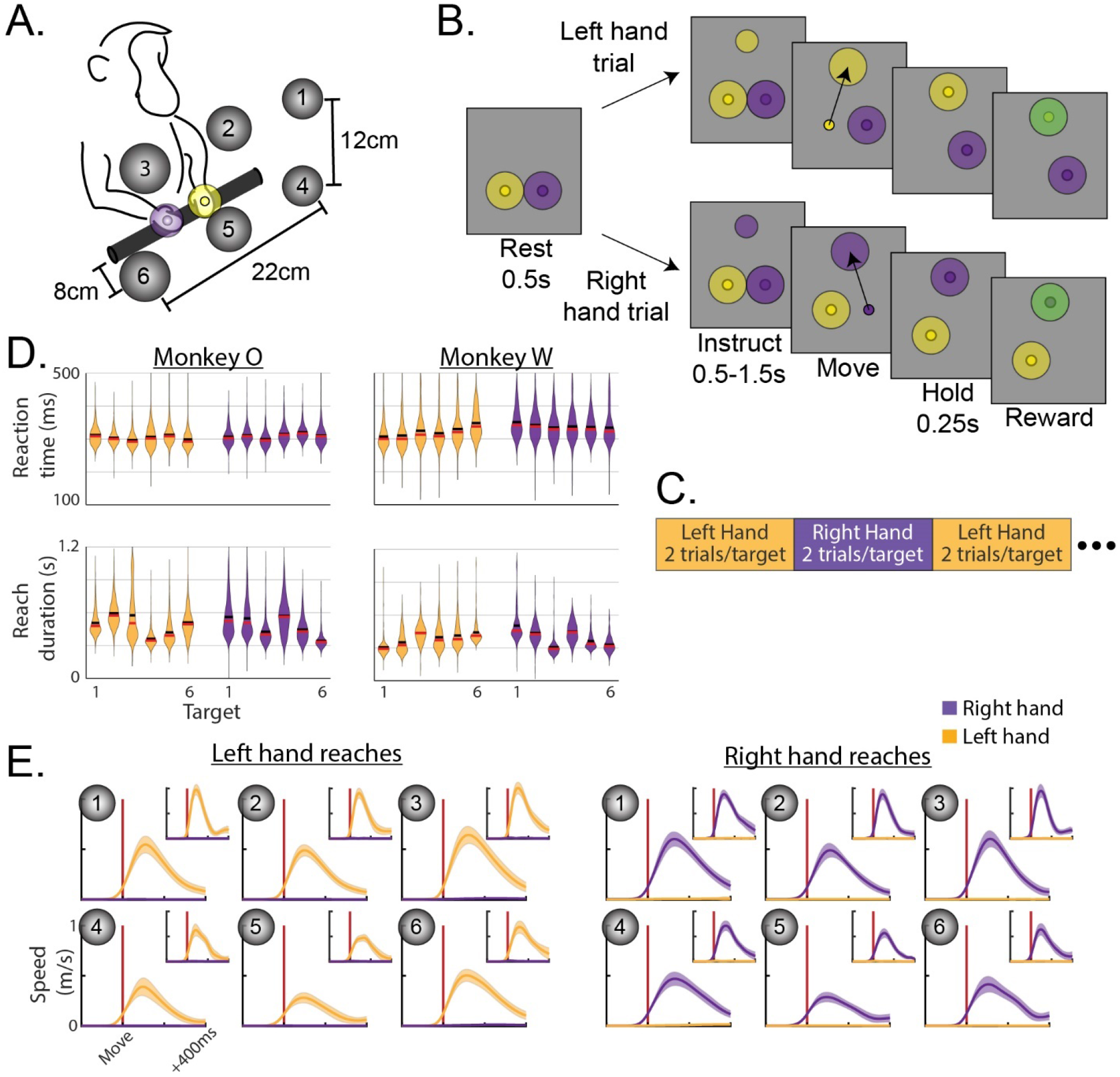
Behavior. (**A**) Monkeys reached to one of six virtual targets, indicated by grey spheres in the cartoon. During the task these would be invisible until one appeared to instruct the reach. (**B**) Trials consisted of 3 phases. Each trial was initiated by placing both hands in start targets and remaining still for 500ms (Rest phase). A small target then appeared at the location of the future reach in a color that indicated which hand to use. The monkey remained still during cue presentation for 0.5-1.5s (Instruct phase). The start target for the reaching hand then disappeared while the reach target enlarged to cue movement (Move phase). (**C**) Hand assignments followed a blocked schedule. (**D**) Distributions of reaction times (top row) and reach durations (bottom row) for each monkey, hand, and target. Left hand reaches in yellow, right in purple. Horizontal black bars show means, red bars show medians. (**E**) Speed profiles during left-or right-hand trials. Both reaching and stationary hands are plotted in each, although stationary speeds are near 0 and hardly visible. Vertical red lines indicate threshold crossing to mark movement onset. Monkey O main, monkey W inset. Mean +/−standard deviation.

Average success rates were above 95% for both hands in both monkeys. Overall, reaction times averaged 308 ms for monkey O and 333 ms for monkey W. Distributions of reaction times for each hand/target combination are displayed in Figure 1D, which were fairly consistent across targets. Reach biomechanics varied across the workspace, resulting in slightly different reach durations across targets (Figure 1D). In terms of kinematics, the initial feed-forward portions of reaches were smooth and stereotyped (Figure 1E). There was a very slight but significant increase in the speed of the non-reaching hand between Rest (mean – monkey O: 1.1 mm/s; monkey W: 2.9 mm/s) and Move (mean – monkey O: 3.6 mm/s; monkey W: 7.6 mm/s) phases of the task (permutation test – monkey O: p=1.0e-4; monkey W: p=1.0e-4). We note that the task was designed to mimic natural reaching without the use of physical restraints. As such, we assume the small movements in the non-reaching arm are part of the normal behavioral repertoire occurring during natural unimanual reaching. Nonetheless, we will address any reasonable impacts these small movements may have in our neural analyses.

### Arm-dedicated units emerge across task phases while the overall distribution remains relatively arm-neutral

We recorded 433 and 113 single-units in the caudal aspect of dorsal premotor cortex (PMd) in monkeys O and W, respectively, and 331 and 289 single-units in primary motor cortex (M1) (Figure 2). Since both arms were used in the behavior, we can evaluate the ipsi-and contralateral responses in each unit. Units were pooled across hemispheres in the analysis, with contralateral summaries reflecting the collection of responses during trials performed with the contralateral arm, and vice-versa for trials performed with the ipsilateral arm. PMd and M1 units were analyzed separately. Firing rates were soft-normalized using the Rest phase mean and standard deviation, and modulation strength is expressed as the mean squared value of these standard scores within the window of interest. This modulation metric is essentially variance, and may be thought of as variance for most purposes.

**Figure 2.**
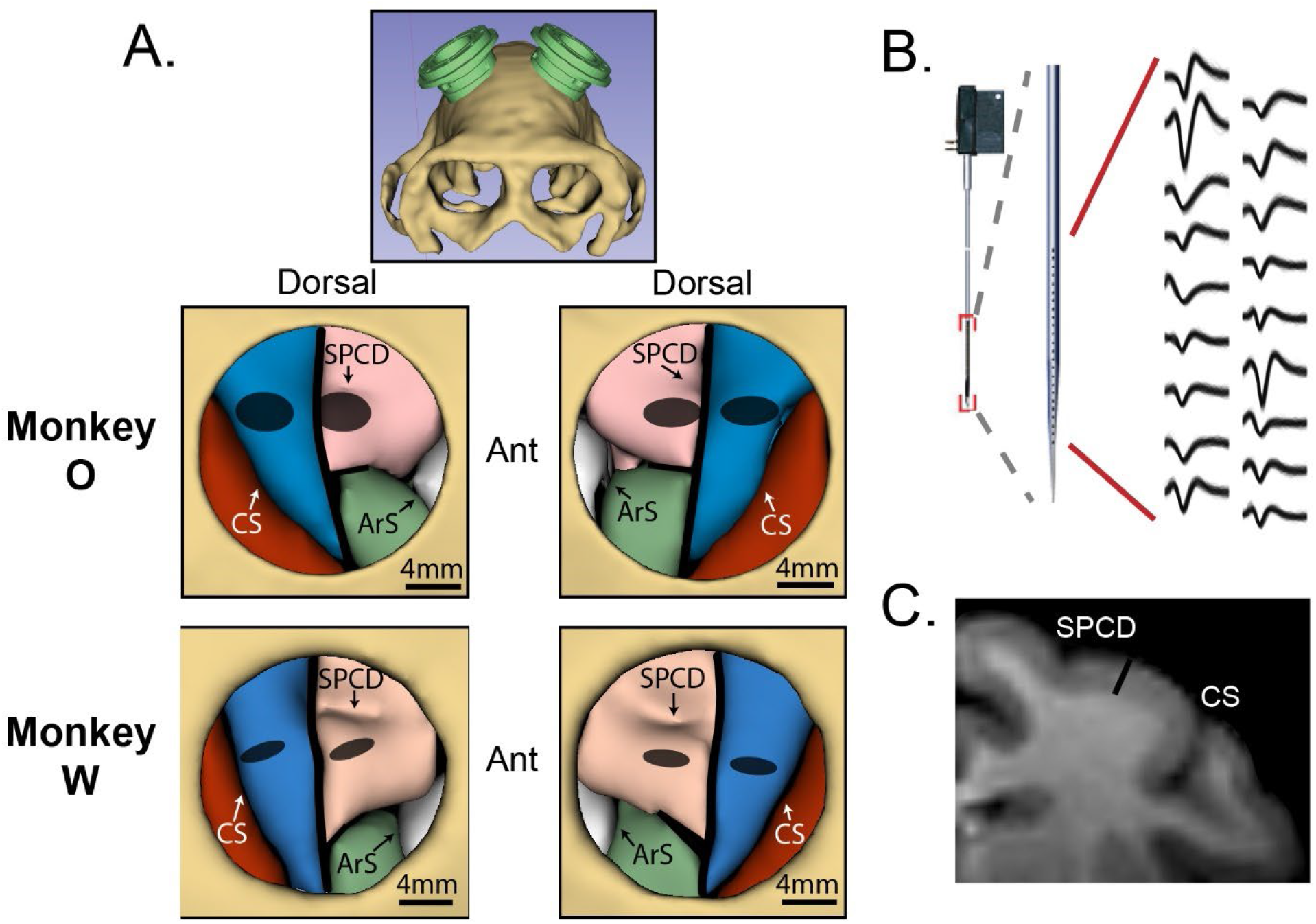
Neural recordings. (**A**) MRI-based volume renderings of the skull and target brain regions. Top panel shows the arrangement of the two chambers. Two bottom rows show segmented brain regions within the cranial window of each chamber, for each monkey. Region boundaries were assigned based on Paxinos et al., 2009. Red -somatosensory cortex; blue - primary motor cortex (M1); pink -dorsal premotor cortex (PMd); green -ventral premotor cortex; white -frontal eye field. CS -central sulcus; SPCD -superior pre-central dimple; ArS - arcuate sulcus. Grey ellipses indicate regions sampled by recordings. (**B**) Interlaminar recordings were obtained using V-and S-probes (Plexon, Inc., Dallas, TX) with 24-32 electrodes aligned perpendicular to the cortical surface. Example waveforms were all simultaneously recorded from a single probe. (**C**) MRI coronal slice, monkey O. 3mm black bar is approximately equal to the distance spanned by electrodes on 32-channel probes. Same landmark labels as in (**A**).

We first analyzed single units to determine the degree of modulation during the Instruct and Move phases of the task (Figure 3). Following instruction, many units in both PMd and M1 became significantly modulated for movements of one or both arms (Table S1). Units in PMd were, on average, more strongly modulated during the Instruct period than those in M1 (Figure 4A; permutation test – monkey O: p=0.012; monkey W: p=3.2e-3). This relationship reversed following movement, with average modulation in M1 becoming stronger than PMd (Figure 4A; permutation test – monkey O: p=2.6e-3; monkey W: p=0.012). These results are in line with the view that PMd plays a privileged role in motor preparation. The distributions of modulation values were heavy-tailed and contained some notably extreme values; however, we chose not to apply any outlier criteria. Controls are performed later in our population-level analyses to ensure that results are representative of trends across the entire population rather than a few extreme units.

**Figure 3.**
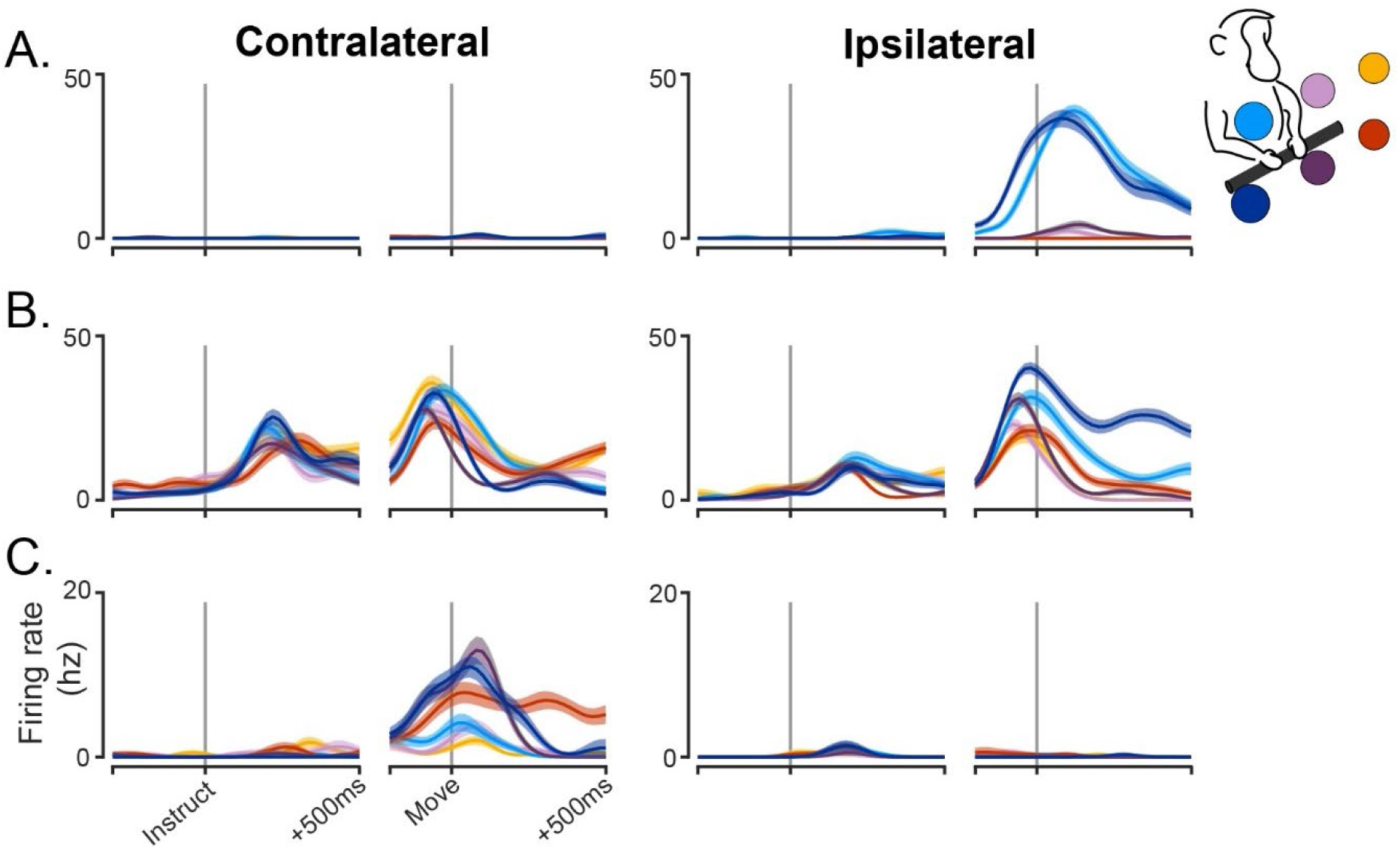
Firing rate traces of example single-units. Trial-averaged firing rates for 3 example single-units, all from the left hemisphere. Each color represents a different target according to the color-coding in the top right. Mean +/−SEM. (A) An M1 unit exclusively modulated during ipsilateral movements. (B) A PMd unit with both Instruct and Move phase modulation for both arms. (C) A PMd unit with modest contralateral modulation during the Instruct phase and strong contralateral modulation during movement, but no modulation on ipsilateral trials.

**Figure 4.**
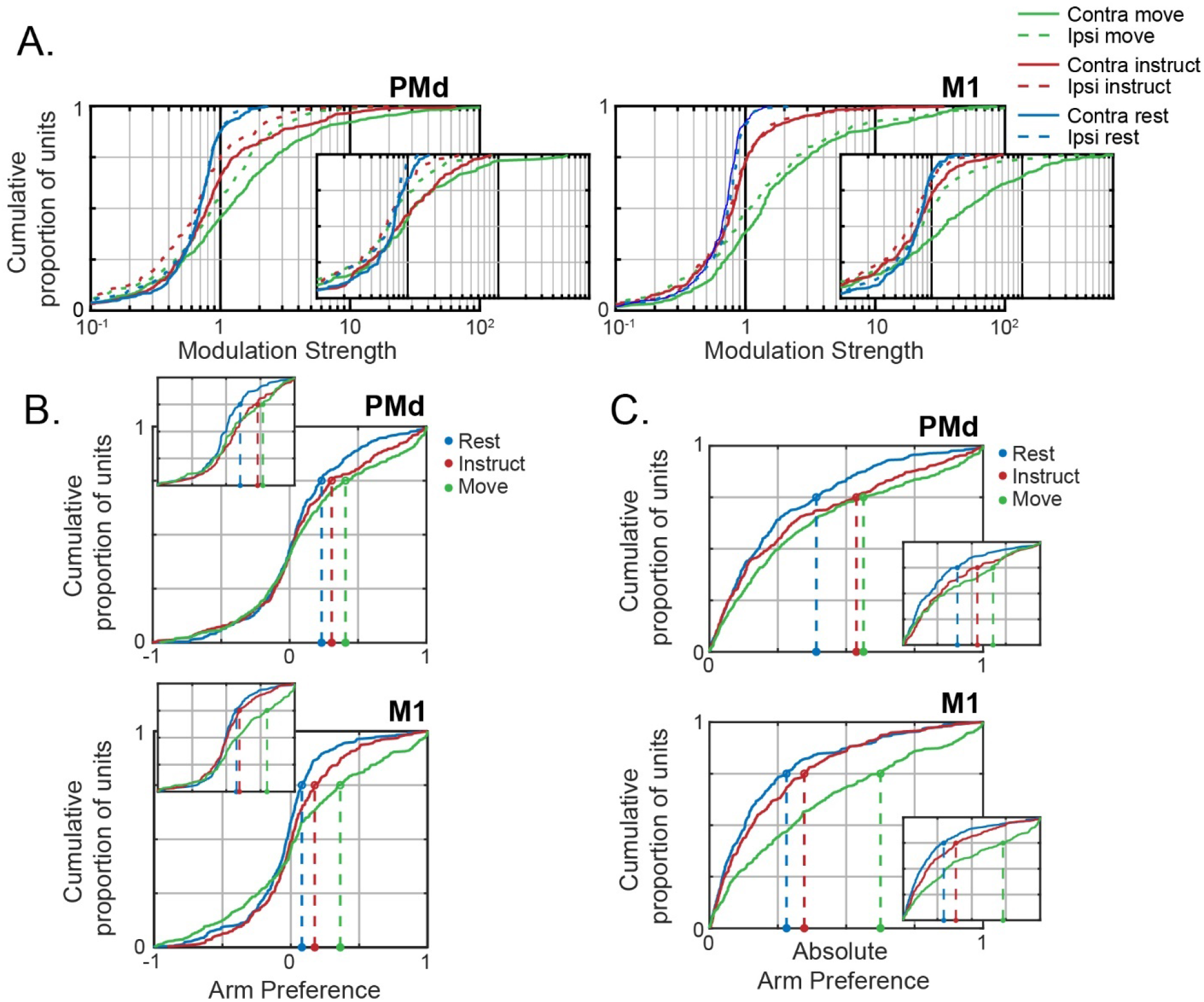
An increasing number of arm-dedicated units emerge with each task phase. (**A**) Cumulative distribution of single-unit modulation during each phase, arm. Left panel PMd, right panel M1. Large values cut off by plot: monkey O Contra Move [134(PMd), 133(PMd), 104(PMd)], Ipsi Move [234(M1), 181(M1), 130(M1)]; monkey W Contra Move [125(M1)]. (**B**) Cumulative distribution of arm preferences during each phase. Top panel PMd, bottom panel M1. Negative values are ipsi-preferring, positive values are contra-preferring. Circles and vertical dashed lines mark the upper quartile of each distribution (**C**) Same as (B), but using the absolute value of arm preference to indicate arm dedication, independent of hemisphere. For all plots: monkey O main, monkey W inset.

We next considered the laterality of each unit by quantifying the relative modulation observed during ipsi-and contralateral trials. We expressed each unit’s arm preference on a scale from −1 to 1, with 1 indicating exclusive contralateral modulation and −1 indicating exclusive ipsilateral modulation (Figure 4B). Although the cue for the forthcoming trial had yet to be presented during the Rest phase, arm selection could be implied from the blocked task structure (Figure 1C). However, except for a very small effect in PMd of monkey O (one-sample t-test – µ_Rest_=0.06, p=9.7e-5), there was no significant contralateral bias observed during the Rest phase in either brain area for both monkeys. Despite the lack of contralateral bias, both monkeys entered arm-specific population states during the Rest phase, which was more pronounced in PMd populations (mean difference between left and right arm firing rates – monkey O PMd: 1.85Hz, M1: 1.64Hz; monkey W PMd: 1.33Hz, M1: 0.98Hz; Figure 4C). For trials in which the same hand was repeated from the previous trial only, it was possible to classify the hand for the forthcoming movement from the population activity (Figure S1).

The emergence of laterality after the onset of the instruction cue mirrored the emergence of general unit modulation: A contralateral bias was present in PMd during the Instruct phase and then became present in both PMd and M1 during movement. Mean arm preference in PMd showed a modest but significant bias in the contralateral direction during the Instruct phase (one-sample t-test – monkey O: µ_Instruct_=0.11, p=7.0e-8; monkey W: µ_Instruct_=0.16, p=1.8e-4) and showed no significant change between Instruct and Move (paired-sample t-test – monkey O: µ_Move_=0.15, p=0.11; monkey W: µ_Move_=0.13, p=0.65). Mean arm preference in M1 did not show a significant contralateral bias until the Move phase (one-sample t-test – monkey O: µ_Instruct_=0.03, p=0.13; µ_Move_=0.07, p=0.013; monkey W: µ_Instruct_=0.02, p=0.31; µ_Move_=0.20, p=5.1e-11).

While shifts in the means were modest, changes in arm preference across phases were most evident in the tails of the distribution, corresponding to units that strongly preferred one arm or the other (Figure 4C). These arm-dedicated units typically preferred the contralateral arm, demonstrated by increased occupancy in the contralateral tails of the arm preference distributions; however, a small proportion of the population was exclusively modulated during ipsilateral trials as well (Figure 4B). Despite much of the population remaining arm-neutral (arm preference near 0) or preferring the ipsilateral arm, the emergence of strongly contra-dedicated units was sufficient to drive contralateral shifts in the population mean. In summary, despite much of the population remaining arm-neutral, an increasing number of highly arm-dedicated units emerged with each task phase, primarily favoring the contralateral arm.

### Modulation preferentially occurs within arm-dedicated units

There are two primary means by which population signals can specify the selected arm at each phase. (1) The population may maintain unique covariance structure for each arm that separates signals along different neural dimensions, even if the constituent units are equally modulated for both arms. (2) Arm-dedicated units may dominate the population response, thereby representing the majority of population variance in dedicated sub-populations. The latter possibility is investigated over the following two sections. First, we consider whether modulation preferentially occurs in units that are strongly dedicated to one arm or the other.

We performed a regression analysis to quantify the relationship between strength of arm preference and modulation for the preferred arm. Importantly, arm preference and modulation were calculated from independent datasets to prevent artificial linkage between the two measures due to sampling noise. A slope of 1 corresponds to an order of magnitude increase in modulation, on average, when comparing fully arm-neutral units with fully arm-dedicated units. As seen in Figure 5A, the slopes are initially near zero and then become positive over time. To quantify these changes, we used a multi-factorial permutation approach to test for effects of Area (PMd, M1), Phase (Rest, Instruct, Move), and Preferred Arm (Ipsi, Contra) on the population slopes.

**Figure 5.**
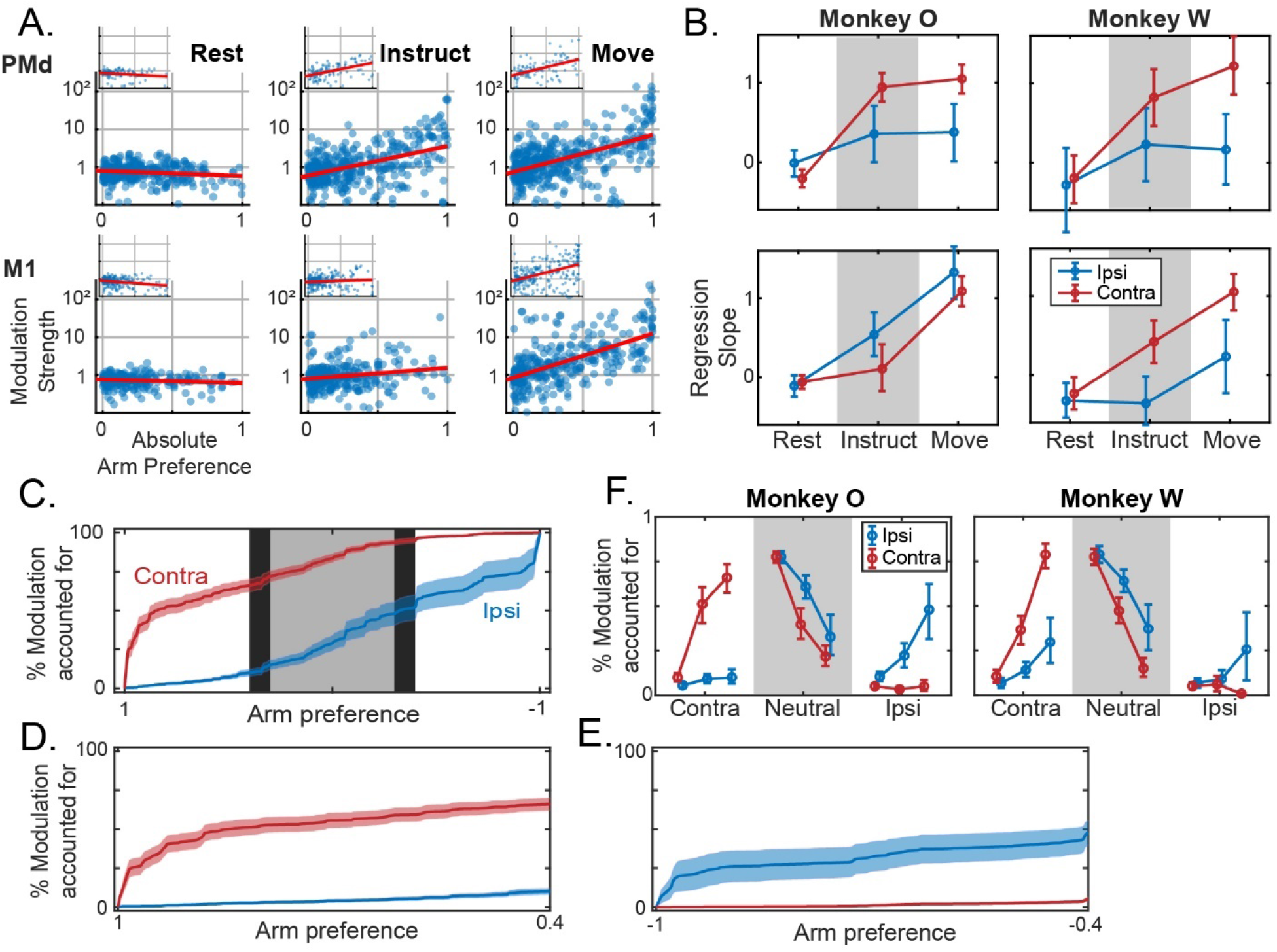
Neural activity is progressively consolidated within arm-specific subpopulations. (**A**) Modulation for the preferred arm plotted against arm preference, for all units in each brain area and task phase. Log-linear best fit lines are displayed in red. Inset figures belong to Monkey W. (**B**) Slopes of regression lines fit to data from (A), independently for ipsi-and contra-preferring sub-populations. Mean +/−bootstrapped 95% confidence interval. (**C-E**) For the Move phase in monkey O, cumulative modulation plotted against arm preference, i.e. each point indicates the proportion of modulation accounted for by all units with arm preference values to the left of the indexed position. Positive values on the x-axis indicate contra-preferring, and negative values indicate ipsi-preferring. Shaded error bars indicate bootstrapped standard error. (**C**) The full spectrum of arm preferences is shown. Shaded backgrounds indicate three partitions: Contra-dedicated [0.4, 1] and Ipsi-dedicated [−1, −0.4] in white, and Neutral [−0.3, 0.3] in grey. (**D**) Cumulative modulation within contra-dedicated regime. (**E**) Same as (D), but ipsi-dedicated. Note inverted axis. (**F**) The proportion of modulation within each partition from (C) during ipsi-or contralateral movements. Note that the total modulation is significantly lower for ipsilateral movements, particularly for Monkey W, and these data are only displayed as proportions. Mean +/−bootstrapped 95% confidence interval.

We found a main effect of Phase in both animals (monkey O: p=1.0e-4, monkey W: p=1.0e-4): a positive correlation between arm preference and modulation strength emerged and strengthened across task phases (Figure 5A-B). By the Move phase, there was approximately a ten-fold increase in the modulation strength of units with an absolute arm preference of 1 (completed dedicated) relative to units with an arm preference near 0 (balanced modulation). Since PMd displayed greater modulation than M1 during preparation but not movement, we tested whether the two areas had differing slopes in each phase independently. We found a significant simple effect of Area during the Instruct phase (monkey O: p=3.0e-4; monkey W: p=6.3e-3) but not the Move phase (monkey O: p=0.13; monkey W: p=0.91). Thus, the relationship was more prominent within PMd prior to movement, while the two areas became roughly equivalent following movement initiation. This was confirmed with a test for 2×2 interaction (monkey O: p=0.025; monkey W: p=9.9e-3). Additionally, we analyzed the relationship between arm preference and modulation for the non-preferred arm to confirm that increased arm preference is associated selectively with increased modulation for the preferred arm (Figure S2). No significant positive relationships were observed in either monkey, either brain area, or any task phase; therefore, greater arm preference is associated with selective increases in modulation for a single arm.

Given the overall contralateral bias, we further tested whether this relationship held for both contra-and ipsi-preferring units. For the contra-preferring units, there was a significant simple effect of Phase (monkey O: p=1.0e-4; monkey W: p=1.0e-4). For the ipsi-preferring units, the Phase effect was significant for monkey O (p=1.0e-4), but only trended in this direction for monkey W (p=0.087), perhaps due to the lesser amount of ipsilateral modulation in monkey W. Slopes were generally steeper for contra-preferring units. The simple effect of Preferred Arm was significant during the Instruct phase for both monkeys (monkey O: p=0.033; monkey W: p=1.0e-4), and significant for Monkey W during the Move phase (monkey O: p=0.53, monkey W: p=1.0e-4). Given that there are also more contra-dedicated units than ipsi-dedicated units, these results suggest that a larger proportion of the contralateral signal exists within dedicated sub-populations compared to the ipsilateral signal. We directly test this conjecture in the following section where we consider population-level implications of these results.

### The population signal is largely confined to arm-specific sub-populations

The preceding analyses establish that there is an increase across task phases in the proportion of units that are strongly dedicated to a single arm, and that those units exhibit greater modulation in activity relative to those that are more neutral. This suggests that the population signal is progressively segregating at the level of individual units. To visualize this segregation, we ordered units based on arm preference and calculated the cumulative modulation at each value, i.e. the proportion of modulation across the entire population that is accounted for by units with arm preferences at or below a certain value (Figure 5C-E). Since PMd and M1 showed similar relationships in the previous analyses, we combined units from the two areas, analyzing them as a collective population. In the extreme case that population signals are entirely segregated, 100% of ipsilateral modulation would occur at an arm preference of −1, and 100% of contralateral modulation would occur at +1.

We focused on two core questions. (1) Does the proportion of dedicated modulation increase across task phases, indicating a progression towards independent signals? (2) Does the amount of independent (or dedicated) modulation differ for ipsi-and contralateral activation? As expected, dedicated regimes of the arm preference distribution captured a large proportion of the modulation associated with movements of one arm and only a small proportion of the modulation associated with the other arm, primarily during execution (Figure 5C-F). For statistical testing, we split the arm preference domain into 3 equal width regimes, corresponding to contra-dedicated (arm preference > 0.4), ipsi-dedicated (arm preference < −0.4), and arm-neutral (−0.3 < arm preference < 0.3) units, and summarized the data by expressing the proportion of modulation contained within each regime (Figure 5F). We again used a multi-factorial permutation approach to test for effects of Phase (Rest, Instruct, Move), and Arm (Ipsi, Contra). We will refer to ipsilateral modulation in the ipsi-dedicated units simply as “ipsi-dedicated modulation” and vice-versa for contra-. We emphasize that these partitions were chosen to broadly isolate the extremes of the distribution; it is not intended that the precise boundaries map onto discrete cell-types or any similar interpretations.

For both animals, the effect of Phase was significant in the contralateral responses (monkey O: p=1.0e-4; monkey W: p=1.0e-4), with the proportion of contra-dedicated modulation increasing across phases (Figure 5F, red lines). Ipsi-dedicated modulation increased across task phases for both monkeys as well (Figure 5F, blue lines), although this effect was only significant for monkey O (p=9.0e-4; monkey W: p=0.31). There was a significant interaction between Arm and Phase for both monkeys (monkey O: p=1.0e-4; monkey W: p=1.0e-4), indicating the stronger emergence of contra-dedicated modulation as compared to ipsi-dedicated modulation. Both animals showed a simple effect of Hand during the Instruct phase (monkey O: p=1.0e-4; monkey W: p=1.0e-4), with more contra-dedicated modulation being observed than ipsi-. This effect was also significant during the Move phase for monkey W (p=1.0e-4) and approached significance for monkey O (p=0.056).

These results suggest that arm signals separate at the level of individual units throughout preparation. Moreover, contralateral signals are more independent than ipsilateral signals, in the sense that a larger proportion of the contralateral modulation was represented in dedicated regimes of the population. Since this characterization of the population response captures most of the modulation for each arm in mutually exclusive sub-populations, we will refer to it as the “dedicated” component.

It is possible, however, that the limited range of movement directions used in the task may influence the degree of dedicated modulation (the maximum angle between target vectors is 107°). For example, a unit that appears dedicated to one arm may only be unmodulated for the other arm over the range of movement directions being tested. That unit may in fact be modulated during different movements, which would cause it to appear neutral if sampled. As a post-hoc control for this possibility, we included data during the return movements following target acquisition, effectively doubling the range of sampled movement directions, and repeated the analyses of Figure 5. The relationship between arm preference and modulation strength remained for this Reach & Return data (Figure S3B-C). This relationship was significant for both monkeys, in both PMd and M1, and for both ipsi-and contra-preferring units, with one exception (permutation test of regression slopes – p<0.05; monkey W, PMd, Ipsi-preferring units p=0.58). The proportion of dedicated modulation was largely unchanged as well (Figure S3D), and replacing Move phase data with Reach & Return did not impact significance of the Phase effect (Contra-dedicated modulation – monkey O: p=1e-4; monkey W: p=1e-4; Ipsi-dedicated modulation – monkey O: p=1.8e-3; monkey W: p=0.22). Therefore, the dedicated signals that we observe persist even with a broad range of movement directions. Returning to the possibilities outlined at the beginning of the previous section, we therefore conclude that this dedicated component progressively becomes the dominant characterization of the population response – dominant in the sense that it represents the majority of modulation across the population.

### Neural subspaces for the two arms diverge across task phases

We next sought to characterize the time course of changes in neural subspaces as movements were prepared and executed. We hypothesized that dedicated activation would drive population signals into diverging subspaces for the two arms. For these analyses, we pooled units from the left and right hemispheres. Using PCA on single-trial data, we first estimated the dimensionality of the neural subspace during each task phase using a cross-validated data reconstruction method (see Methods; Yu et al., 2009). This is an essential step to avoid drawing conclusions based on noise-dominated dimensions. Dimensionality was calculated separately for each session and arm. During Rest, the dimensionality was approximately 5-6, and decreased to approximately 4 during the Instruct and Move phases (Figure 6A). The high dimensionality during Rest aligns with recent reports (Dabrowska et al., 2020), and values during Instruct and Move were comparable to those found in previous studies using similar methods (Yu et al., 2009). We therefore chose to focus on only four components to represent the neural subspaces of each dataset.

**Figure 6.**
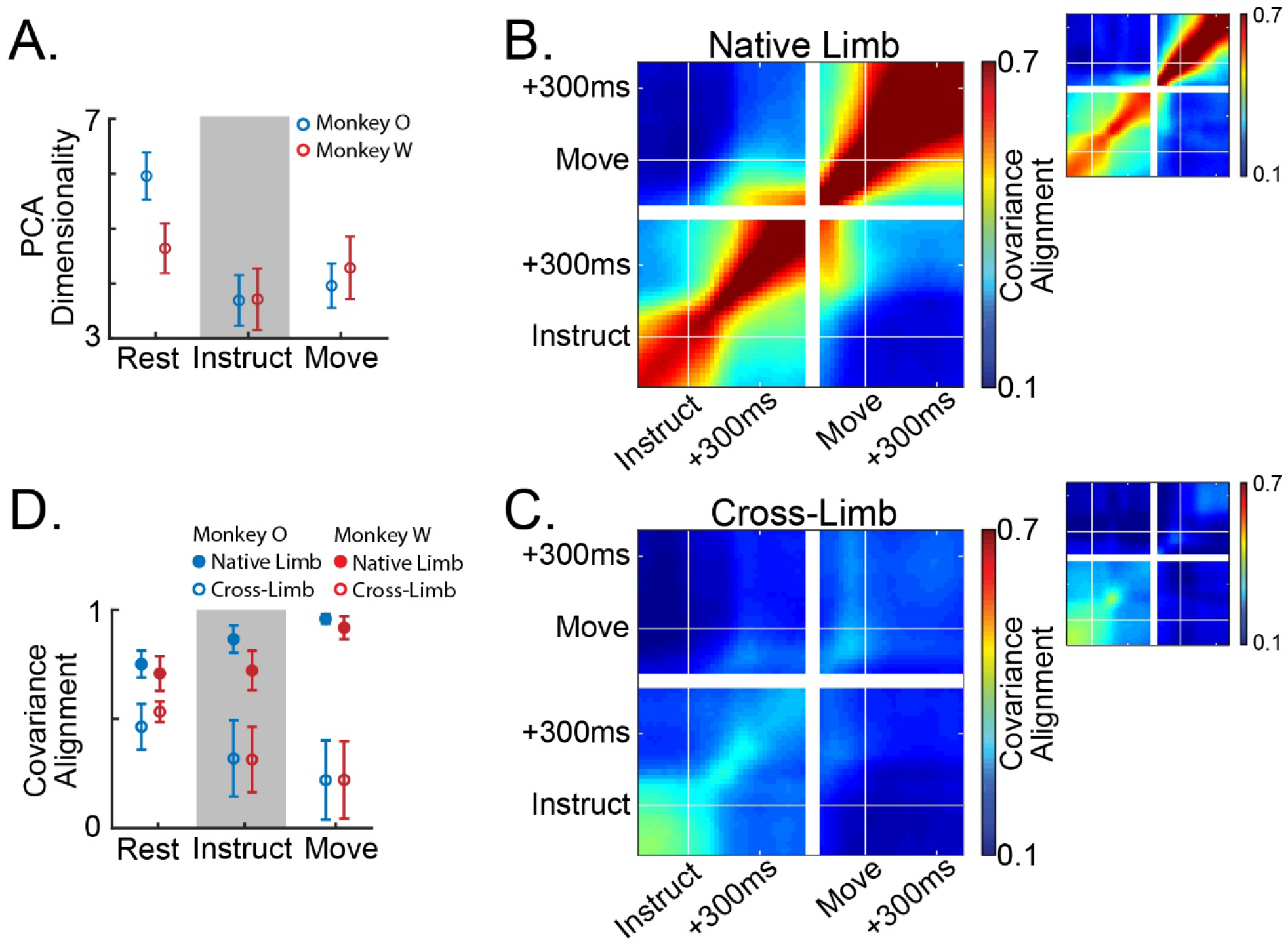
Population activity reorganizes and diverges for the two limbs throughout planning. (**A**) Dimensionality of the PCA subspace estimated as the number of components that minimizes the cross-validated reconstruction error of the full-dimensional neural data. Mean +/−standard error across datasets. (**B**,**C**) Heat maps indicate alignment of 4-dimensional PCA subspaces between all pairs of timepoints across the Instruct and Move phases of the task, averaged across sessions. (**B**) Compares subspaces across time for movements of the same arm. Three blocks forming along the diagonal indicate three distinct subspaces: a pre-instruction Rest space, a post-instruction Instruct space, and a peri-movement Move space. (**C**) Compares subspaces across time for movements of opposite arms. Prior to instruction there is a moderate alignment of the subspaces for each limb, however, the two subspaces diverge around 100ms post instruction. (**D**) Summary of the data in (B,C). Mean +/ − standard deviation across datasets.

We calculated the alignment between PCA subspaces associated with left or right arm movements using a metric that describes the proportion of low-dimensional variance for one dataset that is captured in the low-dimensional space of another (see Methods; Athalye et al., 2017). If the network is organizing activity in the same way across datasets, then the covariance alignment is 1, regardless of signal magnitude. If activity is reorganized into orthogonal subspaces across datasets, then the covariance alignment is 0. Two types of alignment measurements were made: (1) Subspaces were fit to random partitions of trials for the same arm – what we will refer to as “native” alignment – giving us an estimate of natural variability in our subspace estimates when compared over the same time window, and describing the evolution of the motor plan when comparing across time windows. (2) Subspaces were fit separately using trials for either arm and compared with each other – what we will refer to as “cross” alignment – describing the divergence of the subspaces for the two arms at each task phase.

Using single-trial activity event-locked to the onset of instruction and movement, we were able to capture the fine-timescale evolution of any emerging or diverging subspaces (Figure 6B-C). When comparing the native alignment across task phases, we observed the emergence of distinct Instruct and Move period subspaces. Figure 6B shows these data displayed as a continuous heat map with block diagonal structure that coincides with the phase transitions. Within each phase native alignment was high, indicating consistent low-dimensional structure in the population activity that was specific to each stage (Figure 6B; Figure 6D filled circles).

As expected, subspaces for the two arms gradually diverged across task phases (Figure 6C; Figure 6D open circles). On the whole, subspaces for the two arms were significantly less aligned than the (cross-validated) comparisons within the same arm (Figure 6D open vs filled circles; two-way ANOVA, ME comparison type – monkey O: p=3.3e-61; monkey W: p=5.2e-29). Interestingly, subspace divergence was already apparent during the Rest phase (paired sample t-test, native-Rest vs cross-Rest – monkey O: p=1.4e-11; monkey W: p=4.7e-7). As mentioned in our analysis of single-unit arm preferences, this is likely due to predictable arm assignments from the blocked task structure (Figure 1C, Figure S1). Cross alignment decreased significantly as the trial unfolded, reaching a minimum during movement (one-way repeated measures ANOVA – monkey O: p=8.0e-7; monkey W: p=2.4e-6). These results map closely onto the progressive segregation of dedicated signals described in the previous section.

### Subspace separation relies upon dedicated signals

Activity within mutually exclusive sub-populations naturally separates into distinct linear subspaces; as such, we can expect some level of subspace separation as a simple result of dedicated variance. However, it is possible that subspace separation could occur within a distributed representation as well (Ames and Churchland 2019; Heming et al, 2019). This question is especially important in considering units that show relatively balanced modulation for the two arms. Even though these units show similar levels of activity during contra-and ipsilateral movement, it is possible that their population-level contributions are different for each, and thus also contribute to subspace separation.

To investigate the extent to which subspace separation relied upon dedicated activation, we analyzed the structure of PCA subspaces via their coefficient weights. Since components of PCA models form an orthogonal basis set, each can be independently analyzed to determine its contribution to subspace divergence. We fit separate PCA models for each arm and task phase and calculated two statistics for each component: (1) To capture the contribution of a given component to subspace separation, we calculated the ratio of variance it captured for the two arms (right/left). (2) To capture the dependence of a given component on arm-dedicated units, we calculated a coefficient-weighted average of the arm preferences for all units (e.g., if non-zero weights were only given to right arm dedicated units, this value would be 1; if weights were evenly distributed across the spectrum of arm-preferences, this value would be 0). A strong relationship between these two metrics would suggest that subspace separation relies upon dedicated activation.

Indeed, this was the case during both the Instruct and Move phases. Figure 7A-C shows a single session example from the Move phase. The top principle components captured a large amount of the variance for the left arm while capturing little variance for the right arm. Components with a variance ratio strongly favoring the left arm almost exclusively weighted units that were themselves highly dedicated to the left arm. The lower components with more balanced variance ratios distributed weights more evenly across the arm preference spectrum. This pattern was evident in each phase throughout recordings from both monkeys. Figure 7D shows the relationship between right/left variance ratio and coefficient-weighted arm preference for the top five principal components of each dataset. Following the instruction cue, components that strongly discriminated between the two limbs (variance ratio far from 1) primarily weighted units that were themselves highly discriminating. This relationship remained strong as the range expanded during the Move phase. The analysis was also repeated using an alternative normalization method to mitigate the effect of highly modulated units. As expected, mitigating the effect of highly modulated units decreased the magnitude of subspace separation while maintaining the relationship between coefficient-weighted arm preference and variance ratio over the reduced range (Figure S4A). This further illustrates the dependence upon highly arm-dedicated, highly modulated units. In summary, these results suggest that the subspace separation described in the previous section relies upon signals that segregate at the level of individual units.

**Figure 7.**
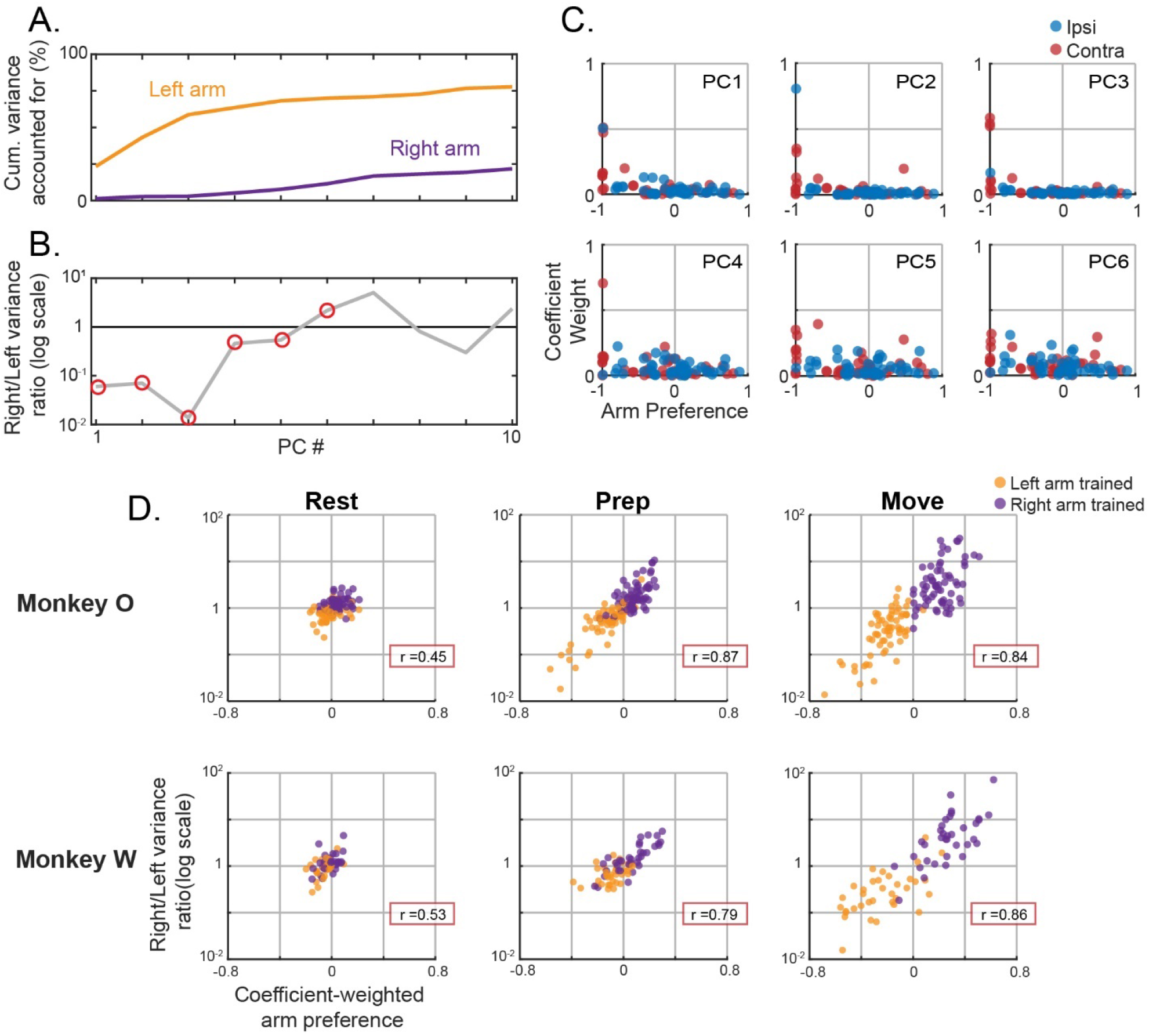
Separation of arm-specific subspaces relies upon unit-level segregation. (**A-C**) Single session example of a PCA model trained to capture bi-hemispheric activity during left arm movements. Held-out testing data for 86 simultaneously recorded units were used. (**A**) Cumulative proportion of variance accounted for across the top 10 principal components. (**B**) For each component, the ratio of the explained variance between the two limbs. (**C**) Absolute values of the coefficient weights for each component plotted against the corresponding unit’s arm preference. Top row represents components 1-3; bottom row represents components 4-6. Positive arm preference values indicate right arm preferring units. (**D**) The component variance ratio for the two arms plotted against a coefficient-weighted average of the arm preferences for each unit in that component. Datapoints represent the top 5 principal components of left or right arm trained models across all sessions. Separate models for each phase are plotted in each column. Pearson correlation coefficient for each dataset is displayed in the red box. Top row monkey O, bottom row monkey W.

### An additional distributed signal contains target-specific information about both arms

The preceding sections make clear that the population signal is dominated by a segregated organization. Nonetheless, it is likely that variance associated with the non-preferred arm of each unit also reflects a meaningful population component, albeit one that is much weaker in magnitude. Indeed, many of the units that we recorded in both PMd and M1 were significantly modulated for both arms throughout preparation and movement (Table S1). To assess the information content and strength of these secondary responses, we divided the entire population of units from both hemispheres and brain areas into two subgroups based on the preferred arm of each unit from a held-out dataset (Figure 8A). If the signals were entirely dedicated to one arm or the other, each subgroup would only contain information about its preferred arm (e.g., a left arm-preferring subgroup would be predictive of left but not right arm movements). If instead there is meaningful activation that is distributed across the same units, then each subgroup would contain both dedicated and distributed information about its preferred arm, but only distributed information about its non-preferred arm.

**Figure 8.**
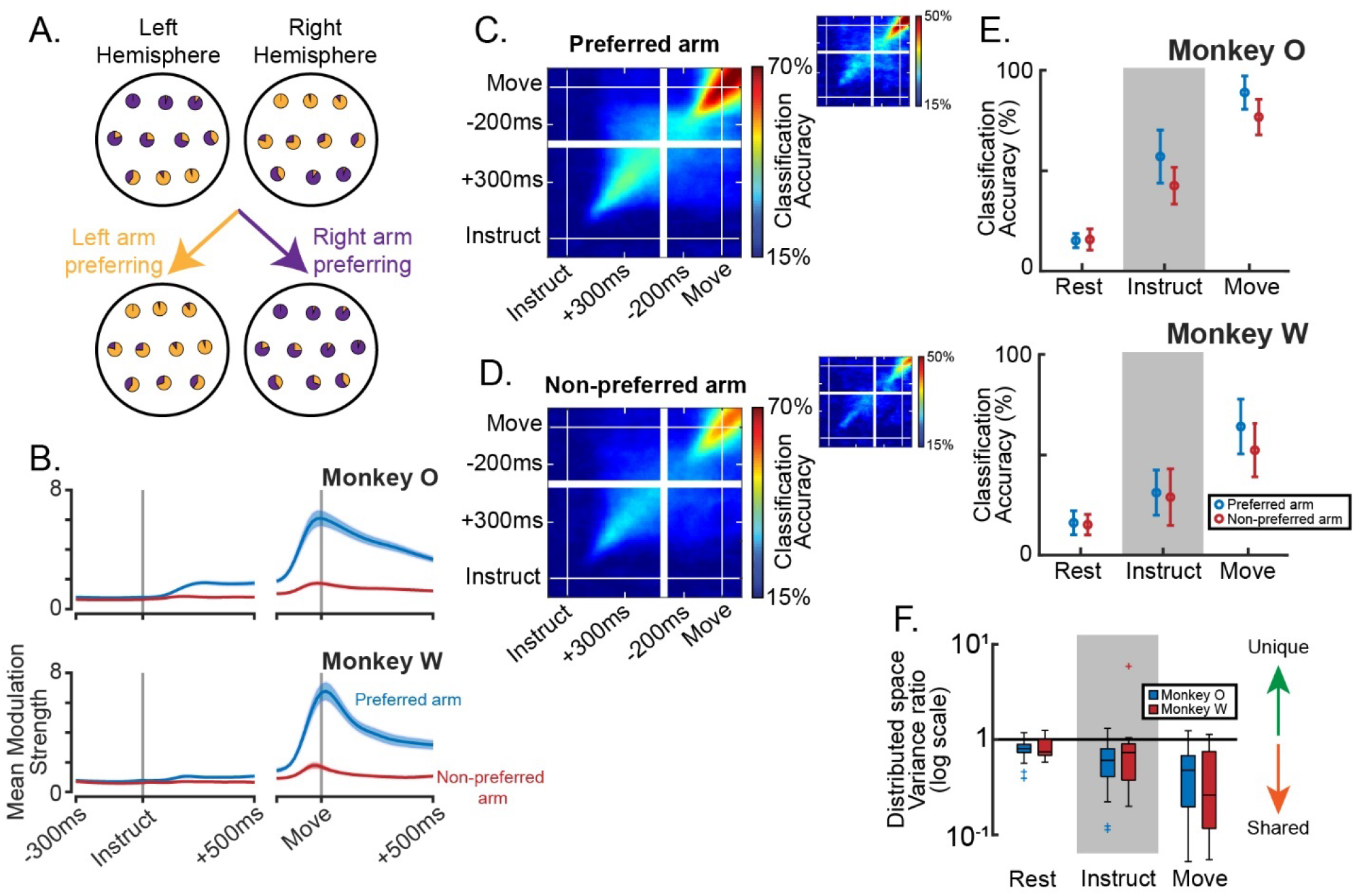
Behaviorally specific information exists within a subspace that captures bilateral activity. (**A**) Illustration of the population partitioning approach. Each unit is represented as a pie-chart displaying the relative modulation during left and right arm trials. Most units in the left hemisphere are more strongly modulated during right arm movements (mostly purple pie-charts), yet some prefer left arm movements (mostly yellow pie-charts). Regardless as to which hemisphere each unit is in, the population may be subdivided into left and right arm preferring sub-populations. On the extreme that all information about each arm is contained within dedicated sub-populations, this simple division will fully segregate the signals such that movements of the non-preferred arm cannot be classified. (**B**) Modulation as a function of time, taken as the mean over all units during trials of their preferred or non-preferred arm, +/−standard error. (**C**) Target classification accuracy using LDA for movements of the preferred arm. Models are trained on each time point and tested on each time point to provide high temporal resolution and inform cross-phase generalization of the classifier. Plots are averaged over all sessions (13 Monkey O, large plots; 7 Monkey W, small plots) and both sub-populations (left-preferring, right-preferring). (**D**) Same as (C), but for non-preferred arm movements. (**E**) Summary data of (C,D) for monkey O, top panel, and monkey W, bottom panel. Mean +/−standard deviation across datasets. (**F**) Ratio of the variance captured in the distributed subspace for the two limbs.

We first analyzed the time course of modulation for each subgroup during movements of the preferred and non-preferred arms. While modulation during preferred-arm trials was much stronger in the Instruct and Move phases, there was a small amount of modulation during trials of the non-preferred arm as well (Figure 8B). To determine whether this modulation carried target-specific information about the behavior, rather than non-specific changes related to task engagement or small movements of the non-selected arm, we trained linear discriminant analysis (LDA) classifiers to predict the target on each trial. Even though the units showed very little modulation when the non-preferred limb was used, prediction accuracy was well above chance (Figure 8C-E, paired sample t-test with Rest – monkey O: Instruct p=1.5e-12, Move p=4.1e-21; monkey W: Instruct p=1.8e-3, Move p=1.1e-7). This suggests that the population code is not entirely dedicated but contains a meaningful distributed component as well. We refer to this as “distributed” in the sense that the contributing units carry information about both arms.

### The distributed signal is contained in a shared subspace for the two arms

We next asked whether subspace separation exists specifically within the distributed portion of population activity. To isolate distributed signals, we again partitioned the population based on preferred arm and fit 4-D PCA models to neural activity during only trials of the non-preferred arm. This is a conservative approach for fitting only the distributed activity, since dedicated activity will be absent during reaches of the non-preferred arm. This approach can also be interpreted as directly isolating the effect of covariance differences by removing the effect of magnitude differences. We will refer to the subspace spanned by these models as the “distributed” subspace.

If population activity for each arm separates along orthogonal neural dimensions, even in the absence of dedicated variance, then the distributed subspace would preferentially capture variance for the non-preferred arm, since that is what it was fit with. Despite having greater magnitude in the ambient space, preferred arm activity would exist largely in the null space of this projection and none of its variance would exist in the distributed subspace. Alternatively, if signals for the non-preferred arm exist within a shared subspace for the two arms, then the patterns of activity for either arm would be preserved through the projection, and we would expect as much or more variance captured for the preferred arm.

Across all task phases and for both animals, more variance was observed in the distributed subspace during preferred arm trials than during non-preferred arm trials (Wilcoxon signed rank – p<0.05 for all six comparisons). The ratios of variance captured for each arm were expressed as non-preferred over preferred and were computed using the raw variance, not the proportion of total variance. Variance ratios were below 1 for nearly every individual dataset and became even lower with each subsequent phase (Figure 8F). Additionally, mean coefficient weights were not significantly different across PMd and M1 for either the Instruct or Move phase, except for the Move phase data in monkey O where M1 weights were slightly larger (permutation test – p>0.05 for monkeys O and W Instruct phase, monkey W Move phase; monkey O Move phase 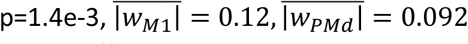). This indicates that the two areas were similarly contributing to the effect. Together these results suggest that across the entire process of preparation and execution of movements, arm signals that are mixed at the level of individual units occupy a shared subspace and are not differentiated through linear population readouts. We again used an alternative firing rate normalization method to confirm that this result was not dependent on overrepresentation of units with the strongest modulation, and the same results were observed (Figure S4B). In summary, the subspace capturing distributed activity is not unique to the arm it was fit to, but rather represents a shared subspace for population activity associated with either arm.

## DISCUSSION

We have shown that the combined population response spanning PMd and M1 across hemispheres contains two primary components with regards to laterality.

The first is characterized by signals that are segregated at the level of individual units, which we call the “dedicated” component. Activity that emerged following the instructional cue was most prominent in PMd and showed a tendency for stronger modulation in units with greater arm preference. This caused the signals for each arm to begin segregating into mutually exclusive sub-populations and occupy divergent low-dimensional subspaces. During the transition to movement, M1 became strongly engaged and segregation of arm signals became even more pronounced.

The second component leveraged signals that were mixed within units, which we call the “distributed” component. We showed that arm signals were not completely segregated by splitting the population in two based on each unit’s preferred arm and analyzing the responses during non-preferred arm trials. Despite being very small in magnitude, these signals contained target-specific information. In contrast to the natural separability of the dedicated component, however, subspaces fit to this activity captured at least as much variance for the other arm during each phase of the task, suggesting a shared subspace for the two arms that persists across preparation and movement.

### Comparison to previous studies of bilateral arm signals in the motor cortex

Our study adds to a growing body of existing work reporting activity related to both arms in the same motor cortical units during either preparation (Tanji et al., 1988; Hoshi and Tanji, 2002; Cisek et al., 2003) or movement (Tanji et al., 1988; Donchin et al., 1998; Kazennikov et al., 1999; Steinberg et al., 2002; Cisek et. al, 2003; Ames and Churchland, 2019; Heming et al., 2019). Two recent studies have addressed the puzzling presence of bilateral activity in M1 during unimanual behavior (Ames and Churchland, 2019; Heming et al., 2019). Despite many units being active for either arm (similar to our own observations in Figure 4B, Table S1), both studies reported separation of population-level arm signals into distinct neural subspaces. Furthermore, they attributed this separation to covariance changes between the two modes that would cause even signals that are mixed within units to contribute to the effect. In the present study, we build upon these observations and reveal an underlying organizational structure that suggests arm signals are not as mixed within units as they would appear based on distributions of single-unit arm preferences. We show that dedicated signals contribute more to the overall population variance and solely account for the presence of arm-specific subspaces. Signals that are mixed within units reflect a different feature of the population. Our example in Figure 5C-E may be compared to Figure 9 of Ames and Churchland, 2019 and Figure 8 of Heming et al., 2019 to demonstrate that segregation of arm signals into distinct neural subspaces likely arises from activation of exclusive sub-populations. Similar to the top principal components of these other studies, dedicated regimes of the population in our study captured large amounts of variance for one arm while capturing very little for the other. Furthermore, we mapped the development of this signal feature across preparation and movement, showing that it begins to emerge even during preparation (Figure 5, 6) and involves both PMd and M1 (Figure 5).

Not only does this clarify the mechanism of signal separation in motor cortical activity, but it acknowledges a critical heterogeneity in the population response. Dedicated signals represent most, but not all, of the population variance. Signals that are mixed within units (distributed signals) reveal a portion of the population activity that is not independent for the two arms. Effector-independent coding has long been appreciated in PMd (Tanji et al., 1988; Cisek et al., 2003). Recently, Willet et al., 2020 identified separate “limb-coding” and “movement-coding” dimensions in neural activity from the hand knob area of human premotor cortex. The “movement-coding” component represented movements for different effectors in the same fashion. This bears similarity to our “distributed” component, although we only show that activity for each arm resides in the same space, not that its relationship to behavior is invariant. Notably, PMd units were not more heavily weighted in our distributed models. This suggests that even in M1, signals for the two arms are not fully independent.

### Progressive segregation of arm-dedicated signals and its functional significance

To our knowledge, this study is the first to compare low-dimensional population structure during preparation of left vs right arm reaching in neurologically-intact subjects (see Willet et al., 2020 for preparation of attempted movements in a human participant with C4 spinal cord injury). It has been proposed that neural subspaces reorganize between preparation and execution of reaching movements (Elsayed et al., 2016), which we observe in our own data (Figure 6B). A principle interest of this study was to determine how the emergence of arm-specific signals maps onto this reorganization process. Since previous work has shown that the transition from preparation to movement coincides with an increased proportion of lateralized units (Cisek et al., 2003; Li et al., 2015; Soma et al., 2017), we expected activity to progressively segregate at the level of individual units, represented primarily in the contralateral hemispheres, as the population reorganizes between task phases. This was indeed the case, and began even during the instruction phase (Figure 5). Careful inspection of model structure revealed that segregation of arm signals at the individual unit level drove separation of arm-specific neural subspaces gradually throughout the trial (Figure 6C-D; Figure 7). This segregation reached its maximum during movement, thus reducing any concern that small movements of the non-selected arm had an impact on our results or conclusions.

Importantly, contralateral signals were more independent than ipsilateral ones; a larger proportion of contralateral modulation occurred in contra-dedicated units than the reverse case for ipsi-(Figure 5). This was not a surprising result, as contralateral bias in the functional organization of motor cortex has been clearly revealed by effects stroke (Hatem et al., 2016), lesion studies (Brinkman and Kuypers, 1973), and cortical stimulation (Penfield and Boldrey, 1937; Alagona et al., 2001; Montgomery et al., 2013). One candidate hypothesis for the presence of ipsilateral activity is that it supplies an independent control signal. There is some evidence that ipsilateral cortex plays an increased role in movement following hemispheric damage (Brinkman and Kuypers, 1973; Hummel and Cohen, 2006; Dancause, 2006; Wilkins et al., 2020), though not necessarily a beneficial or compensatory one. The magnitude of ipsilateral encoding increases with the degree of movement complexity (Verstynen et al., 2005) and may involve spatially distinct neural populations (Ziemann et al., 1999; Chen et al., 2003). However, the corticospinal tract (CST) is almost entirely contralateral, and the effectiveness of the ipsilateral component has been debated (Lacroix et al., 2004; Rosenzweig et al., 2009; Soteropoulos et al., 2011; Baker et al., 2015). Ipsilateral cortex may also exert its influence via connections made in the reticular formation (Alagona et al., 2001; Baker et al., 2015; Wilkins et al., 2020), which projects bilaterally to the spinal cord. Our results showed a small amount of independent ipsilateral activity (monkey O more so than monkey W), with more of the ipsilateral signal coming from non-dedicated units (Figure 5). Thus, if the ipsilateral hemisphere provides any independent control signal, it is much weaker than the contralateral signal. Rather, our results suggest that ipsilateral signals are involved in some form of bilateral control, which we now discuss.

### Bilateral signals and their role in motor control

Correlated activity for movements of the two arms has been widely reported in the literature, primarily using macro-scale neurophysiological approaches. Increases in excitability of homologous effectors during transcranial magnetic stimulation (TMS) (McMilan et al., 2006) and symmetric activation patterns in functional magnetic resonance imaging (fMRI) (Verstynen and Ivry, 2011; Diedrichsen et al., 2013) suggest that bilateral motor cortical circuits are organized with mirrored properties. Similar correlated structure has also been reported in human ECoG (Bundy et al., 2018) and premotor spiking activity (Willet et al., 2020). Mirror activation and other forms of interhemispheric communication have been proposed to support intermanual skill transfer (Diedrichsen et al., 2013) or shaping of contralateral activity patterns during complex behavior (Verstynen et al., 2005). However, correlations between the tuning for ipsilateral and contralateral arm movements in M1 units tend to be weak or absent (Steinberg et al., 2002; Cisek et al., 2003; Heming et al., 2019). In the present study we have not directly compared directional tuning, yet we did observe that the distributed component of bilateral signals existed within a shared subspace for the two arms (Figure 8). Mirror activity would necessarily reside in the same neural subspace for each arm, provided that subspace is linear, as all linear subspaces are invariant with respect to reflection. Our results are therefore consistent with functional hypotheses of ipsilateral cortex involving mirror symmetric activation, and more generally for hypotheses that predict linear correlations between activation patterns for the two arms. We note, however, that while consistent linear correlations in the tuning properties of individual neurons would deterministically result in shared neural subspaces, a lack of linear correlation does not mean that neural subspaces will be orthogonal.

The distributed component that we have characterized may also play a role in bimanual coordination. Distinct bimanual activity patterns have been observed in caudal premotor regions using human fMRI (Diedrichsen et al., 2013) and in M1 using single-unit recordings in monkeys (Donchin et al., 1998; Steinberg et al., 2002). Surgical transection of the corpus callosum, the primary direct connection between hemispheres (Gazzaniga, 1989), disrupts typical spatial coupling and continuous synchronization of arm movements as well (Franz et al., 1996; Kennerly et al., 2002), suggesting a cortical locus for these forms of bilateral control. These studies suggest that bilaterally distributed networks involving PMd/M1 may facilitate bimanual coordination, a function historically attributed to the supplementary motor area (Brinkman, 1981). Our task involved unimanual movements, containing no component of coordination. However, the result that target-specific information existed within a shared subspace (Figure 8) is consistent with a role in coordination. We make limited claims on this hypothesis due to our simplified behavior, and stress that implicating a role in bimanual coordination does not simply mean revealing a shared substrate for signals of both limbs. Nonetheless, a bi-hemispheric network structure may underly computations for controlling the two arms as a unified plant (Welford, 1968). M1 has been implicated in multi-joint integration for voluntary movement and feedback control (Scott, 2003; Pruszynski et al., 2011). Bimanual behaviors have a similar task of overcoming redundant degrees of freedom (Bernstein, 1967); many patterns of behavior for each arm independently may help one achieve an action goal so long as cooperation of the two remains intact (“motor equivalence”, Lashley, 1933). This lower-dimensional behavioral coordination space, sometimes called “the uncontrolled manifold” (Scholz and Schoner, 1999), would likely have a similar neural manifold in which bilateral arm signals interact (for related discussion and review, see Swinnen and Wenderoth, 2004; Wiesendanger and Serrien, 2004; Diedrichsen et al., 2010). The distributed space that we report may reflect such a manifold.

### Interpretations from a dynamical systems perspective

One unified explanation for the two components identified in this study is that they represent the computational (or “hidden”) layers and the output layer of cortical processing. In this framework, the distributed signal would reflect a bilateral network that plays a supportive role in motor processing rather than direct output. The idea that bilaterally distributed networks contribute to computations that do not directly represent the output has been previously proposed by Ames and Churchland, 2019. Preparatory activity in motor areas reflects abstract features of action and may lack a strong contralateral bias (Hoshi and Tanji, 2002; Cisek et al., 2003). The distinctive lack of laterality in the distributed signal we observed is consistent with other reports of abstract preparatory responses. It played a relatively stronger role during preparation as well, since the dedicated component did not fully develop until movement. This aligns with reports that behaviorally specific features become more apparent in motor cortical signals during active behavior, including laterality (Shen and Alexander, 1997; Cisek et al., 2003).

From a dynamical systems perspective, distributed signals could serve to enforce internal dynamics of the overall population. Preparatory signals in pre-and primary motor cortex are thought to converge on an ideal population state, or initial condition, such that internal circuit dynamics will guide appropriate patterns of activity for the upcoming movement (Churchland et al., 2006; Shenoy et al., 2013; Li et al., 2016). Rodent studies have shown that preparatory activity in motor cortical neurons projecting to other cortical areas lacks strong laterality, while neurons with descending output exhibit pronounced contralateral bias and became active closer to movement onset (Li et al., 2015, left vs right directional licking task; Soma et al., 2017, left vs right arm pedal pressing task). Furthermore, these bilaterally distributed networks provide robustness to unilateral perturbation during preparation, and it has been hypothesized that the two hemispheres operate together to maintain the network state (Li et al., 2016, left vs right directional licking task). The two components that we have identified generally align with this form of network structure. In addition to setting the initial state, persistence of the distributed component during movement may reflect the ongoing dynamics of pattern generation (Shenoy et al., 2013; Sussillo et al., 2015).

Within this interpretation, progressive segregation of arm signals may reflect emergence of descending output from the network that mirrors the well-established laterality of anatomical pathways (Brinkman and Kuypers, 1973; Soteropoulos et al., 2011). Alternatively, it may reflect a timing signal for triggering action or transitioning the network from preparation to movement (Sussillo et al., 2015; Kaufman et al., 2016) while simultaneously specifying the selected effector. Like the dedicated signals we observed, signals that reflect the timing of movements, but not their direction, have been shown to capture the most variance in PMd/M1 population responses (Kaufman et al., 2016). Premotor activity has also been shown to contain “limb-coding” dimensions that specify a movement effector independently of the movement type (Willet et al., 2020). The large dedicated signals that we observe bear similarity to both of these previously identified response features, and all three could reflect the same underlying computational process.

In summary, we present a statistical description of arm signals spanning M1 and PMd throughout reach preparation, characterizing in detail both lateralized and non-lateralized features of the population response. The two components that we have identified will be crucial for contextualizing current theory on bilateral motor cortical processing as well as designing future experiments that investigate the independence and interaction of signals across the hemispheres.

## METHODS

### Behavioral recordings and task

Kinematic data were collected using LED-based motion tracking of several points along each arm (Phasespace Inc, San Leandro, CA). 3D positions of each LED were sampled at 240Hz. Prior to offline analysis, these positions were smoothed using a cubic spline and smoothing parameter 0.005 (*cspaps* function – MATLAB). The most distal LED, located on the back side of each hand just below the wrist, was used for online endpoint feedback and all offline analysis.

Monkeys were trained to perform a variant of an instructed-delay reaching task (Figure 1B). Endpoint feedback of each arm and all visual stimuli were presented to the animal using a custom-built virtual reality 3D display. This display consisted of two mirrors that projected shifted images independently to each eye to produce stereopsis. Cursors, indicating effector endpoint position, were color coded for the left (yellow) and right (purple) hands, as were all associated stimuli.

Each trial began with the appearance of the start positions for each hand (spherical targets, radius 4cm, with centroids separated by 8cm), located near the body on top of a physical bar that the monkey rested its hands on (Figure 1A). In a self-initiated manner, the monkey would assume the start position by placing both cursors in their appropriate starting positions and maintaining that position for 500 ms (“Rest” phase). Our threshold for detecting movement online was 9cm/s; breaking this threshold would abort the trial.

Marking the beginning of the “Instruct” phase, a cue (spherical target, radius 3cm) would appear at one of six locations within a fronto-parallel plane 8cm in front of the start positions (Figure 1A). The color of the cue indicated the required arm, and position of the cue was the target location for the forthcoming reach. The instruction cue remained visible through the delay period, a duration that was sampled uniformly on the interval 500-1500ms. Movement beyond the speed threshold with either hand would abort the trial.

At the end of this period, two simultaneous changes signaled the monkey to move and marked the start of the “Move” phase. First, the sphere defining the start position for the cued arm disappeared. Second, the cue at the target location enlarged (3cm to 4cm radius). The monkey then reached toward the target and once at the terminal location, had to maintain that position for 250ms. To earn a juice reward, the animal had to initiate the reach within 500ms of the onset of the imperative, terminate the movement within the target’s circumference, and keep the non-reaching hand stationary for the duration of the trial. To further emphasize that the trial was successful, the target turned green.

300ms windows were used to represent each phase in data analysis. For the Rest phase, we used the final 300ms before the onset of the instruction cue. For the Instruct phase, we used data in the interval between 200ms to 500ms post-cue. For the Move phase, we used the first 300ms following the onset of movement, defined as when speed of the reaching hand exceeded 10cm/s. We used a late window for the Rest phase to avoid any residual activity associated with moving to the start positions. The steady state neural response was used to position the Instruct phase window; this was reached approximately 200ms after the onset of the instruction cue (see Figure 7B). The Move window was selected to capture peak neural activity associated with movement while including only the feed-forward portion, which typically lasted 250-300ms (Figure 1C, bottom row). Reach durations were calculated as the time between movement onset and the first point where (1) movement speed dropped below 20cm/s, and velocity in the depth direction reached 0.

### Surgical implantation

All procedures were conducted in compliance with the National Institutes of Health Guide for the Care and Use of Laboratory Animals and were approved by the University of California at Berkeley Institutional Animal Care and Use Committee under protocol ID AUP-2014-09-6720-1. Two adult male rhesus monkeys (Macaca mulatta) were implanted bilaterally with custom acute recording chambers (Grey Matter Research LLC, Bozeman, MT). Partial craniotomies within the chambers allowed access to the arm regions of dorsal premotor (PMd) and primary motor (M1) cortices in both hemispheres. Localization of target areas was performed using stereotactically aligned structural MRI collected just prior to implantation, alongside a neuroanatomical atlas of the rhesus brain (Paxinos et al, 2000).

### Electrophysiology

Unit activity was collected using 24-32 channel multi-site probes (V-probe -Plexon Inc, Dallas, TX), with contacts separated by 100um and positioned axially along a single shank. Probes were lowered deep enough to cover roughly the full laminar structure of cortex (Figure 2B-C). The depth of insertion was determined by (1) measurements of the dural surface prior to recording, and (2) presence of spiking activity across all channels. 2 probes were typically inserted in each hemisphere daily and removed at the end of the session, one in PMd and one in M1. A total of 12 insertion points across PMd and M1 of each hemisphere were used across 13 recording sessions in Monkey O, and 6 insertion points across 7 sessions for Monkey W (Figure 2A).

Neural data were recorded using the OmniPlex Neural Recording Data Acquisition System (Plexon Inc, Dallas, TX). Spike sorting was performed offline (Offline Sorter – Plexon Inc, Dallas, TX). Single-unit waveforms were isolated in multi-dimensional feature space (including principal components, non-linear energy, waveform amplitudes) and rejected if either (1) the waveform clusters were not stable over the course of the session, or (2) >0.4% of inter-spike-intervals were below 1ms. For population level analyses (PCA, LDA), a small number of multi-units were included. A multi-unit was defined by waveform clusters that separated from the noise cluster and were stable over time, but did not quite meet the inter-spike-interval criteria or contained what might be multiple unit clusters that could not be easily separated. For monkey O, the average proportion of multi-units in each single session population sample was 17%, ranging 12-25%. For monkey W, average 20%, ranging 12-32%.

Spiking data were binned in 20ms non-overlapping bins, square-root transformed to stabilize variance, and smoothed with a 50ms gaussian kernel for all analyses (Yu et al., 2009).

### Modulation and Arm Preference metrics

As a time-varying value, modulation was calculated as:

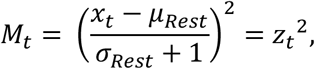

Where

*x*_*t*_: *instantanesous firing rate at time t*

*μ* _*Rest*_: *mean firing rate during Rest*

*σ* _*Rest*_: *standard deviation during rest*

This unitless metric reflects the deviation from baseline activity, normalized by baseline fluctuations. The constant 1 was added to the denominator for soft-normalization to ensure that units which were silent during rest did not have exploding values and were not overly emphasized in the dataset. Because some units had slightly different activity on left and right arm trials even before instruction, the standard deviation during Rest was calculated separately for each arm and *σ*_*rest*_ was calculated as the mean of the two.

Single values of modulation representing each phase were obtained using the same 300ms phase windows that were used in behavioral analysis (see Methods – Behavioral recordings and task). Phase-specific modulation data were concatenated across trials into a (16 *m x*1) vector, where *m* is the number of trials and 16 is the number of samples within a 300ms phase window. The mean over this vector provided our scalar estimate of modulation. Note that this is simply a normalized form of variance, which makes the comparison of single-unit and population-level results more straightforward. To make the relationship with variance explicit, modulation can be rewritten as:

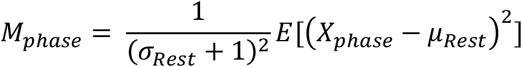

Where the expectation on the right is essentially variance using the Rest mean.

Arm Preference was calculated independently for each phase of the task using the formula:

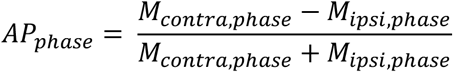

An arm preference of 1 corresponds to a unit that is exclusively modulated during contralateral trials, while an arm preference of −1 is the same for ipsilateral trials. In Figure 7, Figure S4, and the accompanying analyses, the convention of left arm and right arm was used in place of ipsi and contra. In analyses that used arm preference along with other features, independent datasets were used to calculate each to avoid any artificial coupling due to sampling noise, e.g. modulation and arm preference. Note also that the scaling factor used in the modulation calculation cancels out of the arm preference calculation, making it invariant to the choice of normalization.

### Principal components analysis

Principal components analysis (PCA) was used to identify low-dimensional representations of population activity with the *pca* function in MATLAB. PCA computes an orthogonal basis set that reflects the principal axes of variation in the data. Individual components do not strictly correspond to observed activity patterns, and one should be wary of interpreting them as such, yet the low-dimensional space spanned by the top few components has been frequently used in systems neuroscience as a helpful descriptor of coordinated ensemble activity (Cunningham and Yu, 2014). PCA was selected over other dimensionality reduction techniques for its widespread use and relative lack of assumptions. Additionally, PCA was used in two recent papers covering similar topics to this one (Ames & Churchland, 2019; Heming et al., 2019). Therefore, using PCA over other alternatives was also intended to improve generalization of our results to the existing literature.

Prior to fitting the models, firing rate data were soft-normalized using the same method as in the modulation strength calculation:

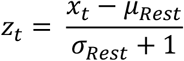

An alternative normalization factor was used to create Figure S4, replacing the denominator by the full firing rate range + 5Hz (Elsayed et al., 2016; Ames and Churchland, 2019; Heming et al., 2019).

Since Rest phase mean activity was already subtracted from individual units, we did not de-mean again prior to computing PCA models. Measures of variance accounted for were not inflated by capturing means because they were computed using the variance of the component scores (Figure 7,8F):

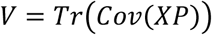

Where *X* is a (16 *m x n*) data matrix of concatenated trials and *P* is an (*n x p*) projection matrix, with *m* trials, 16 samples in the phase window of each trial, *n* units, and *p* principal component dimensions.

Cross-validation approaches were used for all analyses and figures to address overfitting. This provided accurate and generalizable estimates of variance capturing metrics that could also be appropriately compared across datasets (i.e. across time or arms).

### Dimensionality estimation

Dimensionality of the PCA subspace was estimated by optimizing the cross-validated reconstruction of full-dimensional neural data from component scores. Given *n* units, *m* trials, and 16 samples within the phase window for each trial, the following procedure was used:

1. Leave out the *i*^th^ trial (16 samples) from the data matrix, yielding training data, *X*^(− *i*)^ ∈ ℝ^16(*m* − 1) *x n*^, and testing data, *X*^(*i*)^∈ ℝ^16^ ^*xn*^.

2. Train PCA model of dimension *p* < *n* on *X*^(− *i*)^, using singular value decomposition (SVD) to compute the projection matrix, *P* ^(− *i*)^∈ ℝ^*n x p*^

3. Leave out the *j*^th^ unit from the testing data and projection matrix by removing the *j*^th^ column and row from each, respectively, yielding 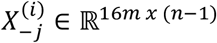 and 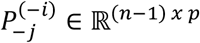. This is the current unit that will be reconstructed.

4. Using the Moore-Penrose pseudoinverse, find a new projection matrix with the *j*^th^ unit removed, whose transpose is 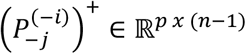. This matrix projects the (*n* − 1) dimensional neural activity into the original *pp* dimensional PC space, therefore computing component scores in the absence of unit *j*.

5. Calculate the component score for the *i*^th^ trial using the remaining units and the new projection matrix, then estimate the *j*^th^ unit from that component score by projecting back into the ambient space. As a single step, this calculation is:

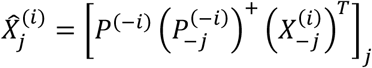

7. Repeat for trials *i*=1,…,*m*

8. Repeat for units *j*=1,…,*n*

9. Repeat for component numbers *p*=1,…,10

10. Take the number of components that minimizes the predicted residual error sum of squares (PRESS) statistic:

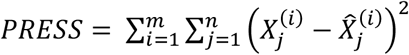

This method reconstructs the full-dimensional neural data, independent of the training set, by identifying consistent population structure. There are no mathematical constraints favoring increased dimensionality, i.e. it is robust to overfitting. As such, the number of components that minimizes the reconstruction error provides a conservative estimate of the dimensions that meaningfully reflect population structure. Similar methods have been used previously for assessing dimensionality reduction techniques for neural data (Yu et al., 2009). Using heuristics, such as the number of components to explain 90% variance, would be inappropriate for our analyses. They are prone to overfitting, which would include meaningless components and impair analysis of model structure via coefficient weights.

### Covariance alignment

We computed a measure of similarity between pairs of subspaces that we call Covariance Alignment. Our method is essentially the same as that previously used for comparing low-dimensional spaces via factor analysis (Athalye et al., 2017). In short, this measure computes the proportion of low-dimensional variance from one dataset that is also captured in the low-dimensional space of another dataset.

Given data matrices *X* _*A*_, *X*_*B*_ ∈ ℝ^*16m x n*^, the following procedure was used:

1. Train PCA models of dimension *p < n* on *X* _*A*_ and *X*_*B*_, using SVD to compute the projection matrices *P*_*A*_, *P*_*B*_ ∈ ℝ ^*n x p*^

2. Project *X _A_* into its own *p*-dimensional space and compute the variance as:

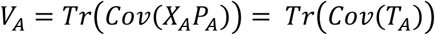

3. Project the *p*-dimensional representation of *X* _*A*_, which is *T*_*A*_, into the *p*-dimensional space identified using *X*_*B*_ and compute the variance as:

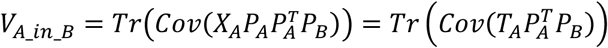

4. Return the proportion of *p*-dimensional variance from dataset *A* that is also captured in dataset *B*’s subspace using the ratio:

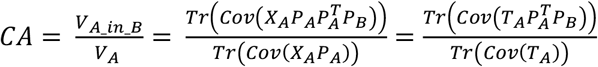

This metric is subtly different from the alignment indices used in Elsayed et al., 2016 and Heming, Cross et al., 2019. The key difference here is the double projection in the numerator, which means that we are specifically capturing the proportion of low-dimensional variance from one dataset that is captured in the low-dimensional space of another, rather than the ratio of overall variance captured in two different subspaces.

### PCA coefficient analysis

Since components of PCA models form an orthogonal basis set, each was independently analyzed to determine its contribution to subspace divergence. Two statistics were calculated for each component using held-out datasets.

First, we projected activity during trials of each arm onto a single component, calculated the variance of the projections for each arm, and expressed them as a ratio. This captured each component’s contribution to discrimination between the arms. For component *C*, this calculation is:

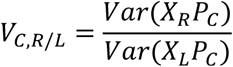

Where *X* _*R*_, *X*_*L*_ ∈ ℝ ^16 *m x n*^ are data matrices for the right and left arms, respectively, and *P*_*C*_ ∈ ℝ^*n x 1*^ is the projection matrix for component *C*. The log of this ratio will be far from 0 if there is much more variance for one arm than the other along the axis defined by *P*_*C*_.

Second, we calculated a coefficient-weighted average of the arm preferences for all units. If non-zero weights were only given to right arm dedicated units, this value would be 1; if weights were evenly distributed across the spectrum of arm-preferences, this value would be 0. Therefore, this measure captured the dependence of a given component on arm-dedicated units. The coefficient-weighted arm preference, *CAP*, for component C was calculated as

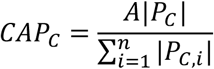

Where *A* ∈ ℝ ^*1 x n*^ is the vector of arm preferences for each unit.

### Linear discriminant analysis

Population coding of movement was analyzed using Linear Discriminant Analysis (LDA) with the *fitdiscr* function in MATLAB. LDA assumes that each class (target x limb combination) is associated with a multivariate normal distribution over the predictor variables (spiking activity of multiple units) having identical covariance but different means.

The feature matrix *X*_*LDA*_ ∈ ℝ^*m x n*^ consisted of a single sample per trial for each of the *n* units. For fine timescale analysis, this was the instantaneous firing rate. For models representing an entire phase, this was the mean firing rate during the 300ms phase window. Uniform priors were enforced for all models. As it was expected that the covariance may change across use of the two arms during reaching, LDA models were trained separately for each limb to allow fitting of arm-specific covariance matrices. LDA was chosen for its robustness to violations of the given assumptions and its history of success with neural data (Diedrichsen et al., 2013; Rich and Wallis, 2016).

### Fine timescale analysis of population coding and subspace development (heatmaps)

The same basic method was used for displaying fine timescale changes in population coding of movements (via LDA) and covariance structure (via PCA, Covariance Alignment). The method is depicted schematically in Figure S5. Neural data were organized as 3D tensors (units, time windows, trials). Comparisons were made between all possible pairs of time windows, using fully independent trial sets to prevent overfitting. For LDA models, this consisted of leave-one-out cross-validation; for Covariance Alignment, random partitioning into two datasets of equal trial numbers. Averages of the cross-validated results provided the 2D matrices visualized using heatmaps in Figure 6B-C and Figure 8C-D. A single row or column therefore reflects the similarity of population coding or covariance between a single timepoint and all other timepoints across the trial. Block diagonal structure in the heatmaps reveals locally consistent structure within task phases.

### Permutation testing procedures

Permutation tests were used for both single and multi-factorial hypothesis testing when parametric tests were inappropriate. Null distributions were constructed by constraining permutations to only data that were exchangeable under the null hypothesis (Anderson and Braak, 2002). For example, we maintained the crossed structure of Phase (Rest, Instruct, Move), by only permuting Phase labels within units. 10,000 permutations were used for all analyses, and p-values were estimated as the proportion of permutations resulting in test statistics that were at least as extreme as what was observed. In cases where the observed test statistic was more extreme than any permutations, we assigned a p-value of 1/number of permutations = 1.0e-4.

## SUPPLEMENTARY FIGURES

**Table S1.**
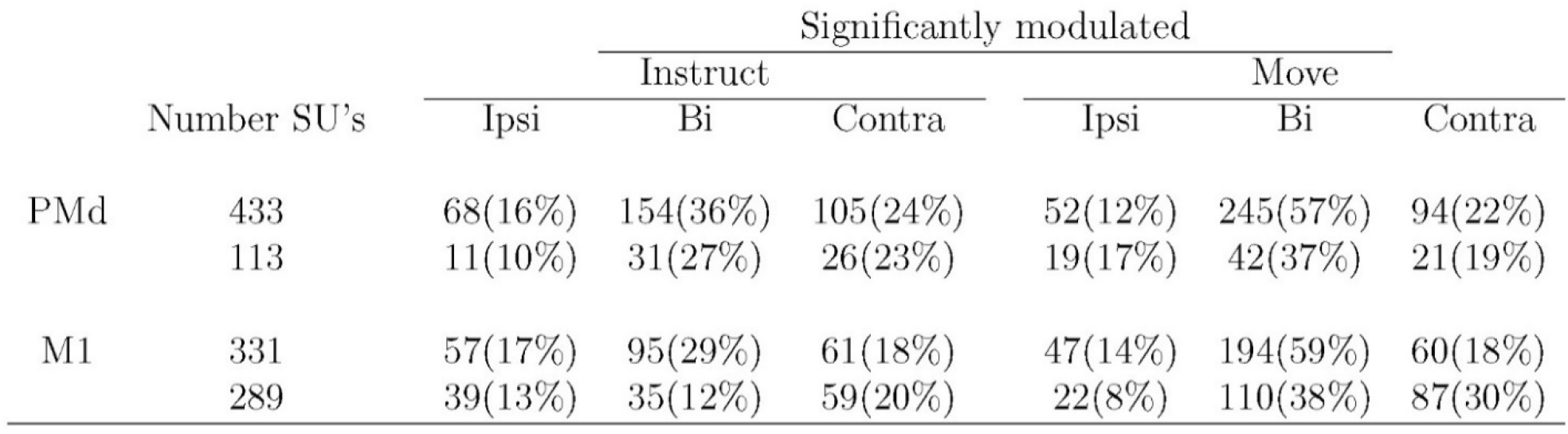
Proportions of significantly modulated single-units across task phases. For well isolated single-units in each brain area, the proportions of the total population that were significantly modulated when compared with the Rest phase (two-sample t-test, p<0.05). For each phase, single-units were classified as uniquely ipsi, contra, or bilaterally modulated. Top row in each pair of rows represents Monkey O, bottom row Monkey W.

**Figure S1.**
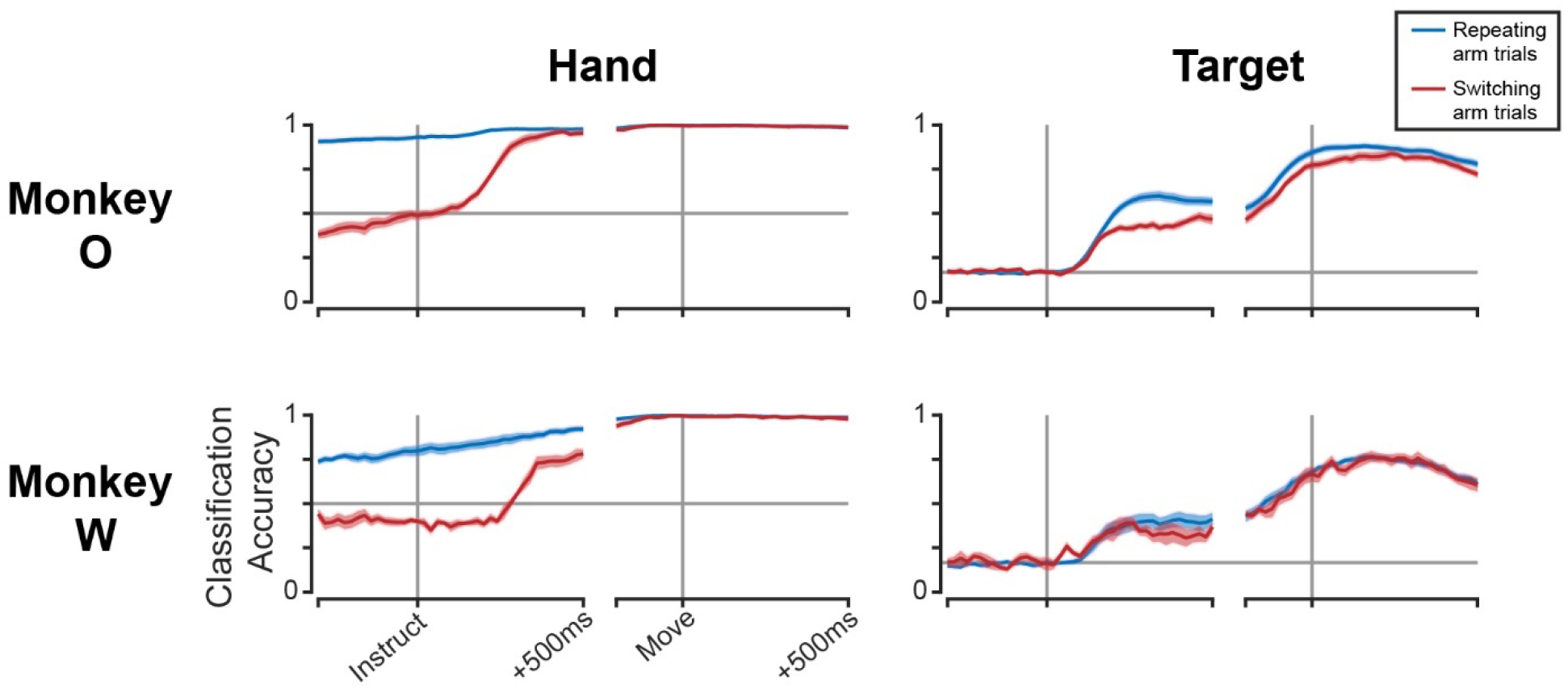
Arm-specific neural patterns exist during Rest on predictable trials. Cross-validated classification accuracy for hand (left column) and target (right column) assignments. LDA models were trained on only trials that required use of the same arm as the previous trial, then tested on either held-out repeating arm trials (blue lines) or switching arm trials (red lines). Separate models were used for each timepoint. Horizontal grey lines indicate chance level. 13 Sessions for monkey O (top row); 7 sessions for monkey W (bottom row). Mean +/− standard error across sessions.

**Figure S2.**
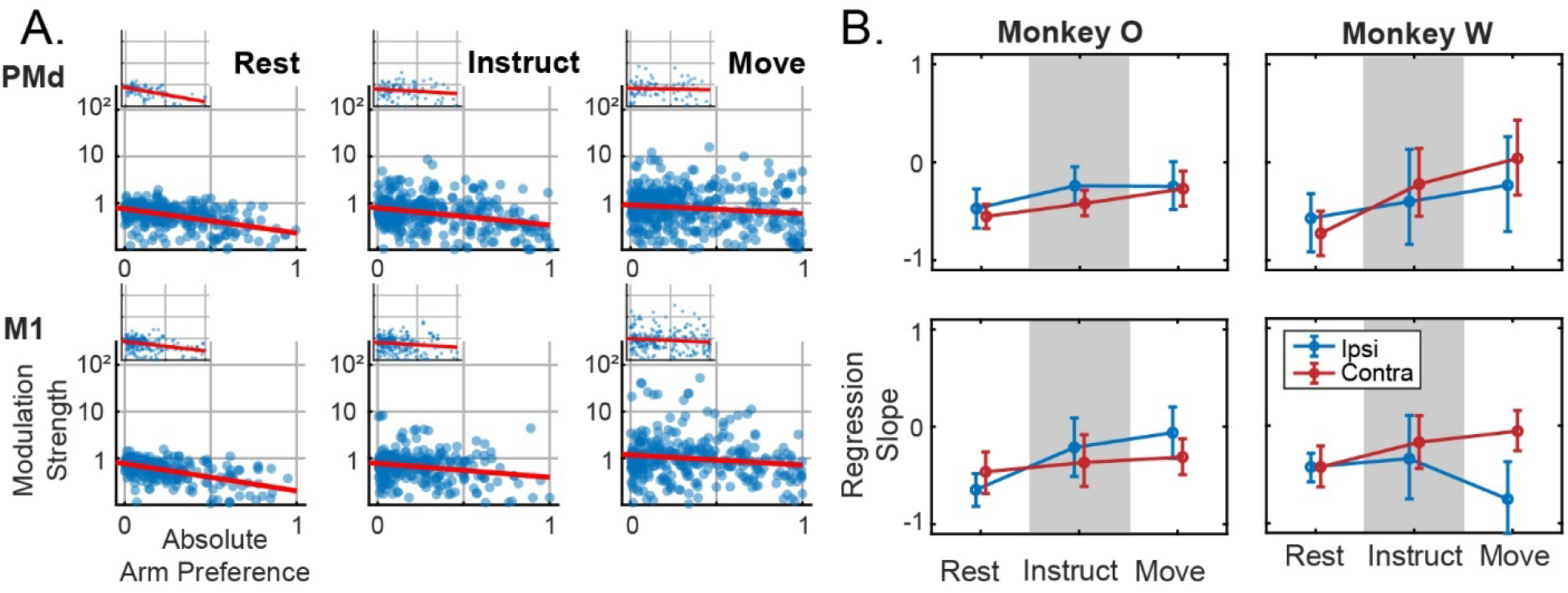
Modulation for the non-preferred arm does not increase with greater arm preference. Companion figure for Figure 5A-B. (**A**) Modulation for the non-preferred arm plotted against arm preference, for all units in each brain area and task phase. Log-linear best fit lines are displayed in red. Inset figures belong to Monkey W. (**B**) Slopes of regression lines fit to data from (A), independently for ipsi-and contra-preferring sub-populations. Mean +/− bootstrapped 95% confidence interval. Note the different y-axis from Figure 4B. The slope was not significantly greater than 0 in any condition, meaning that increased arm preference is associated uniquely with greater modulation in the preferred arm, as opposed to an increase for both arms that is just larger for the preferred arm.

**Figure S3.**
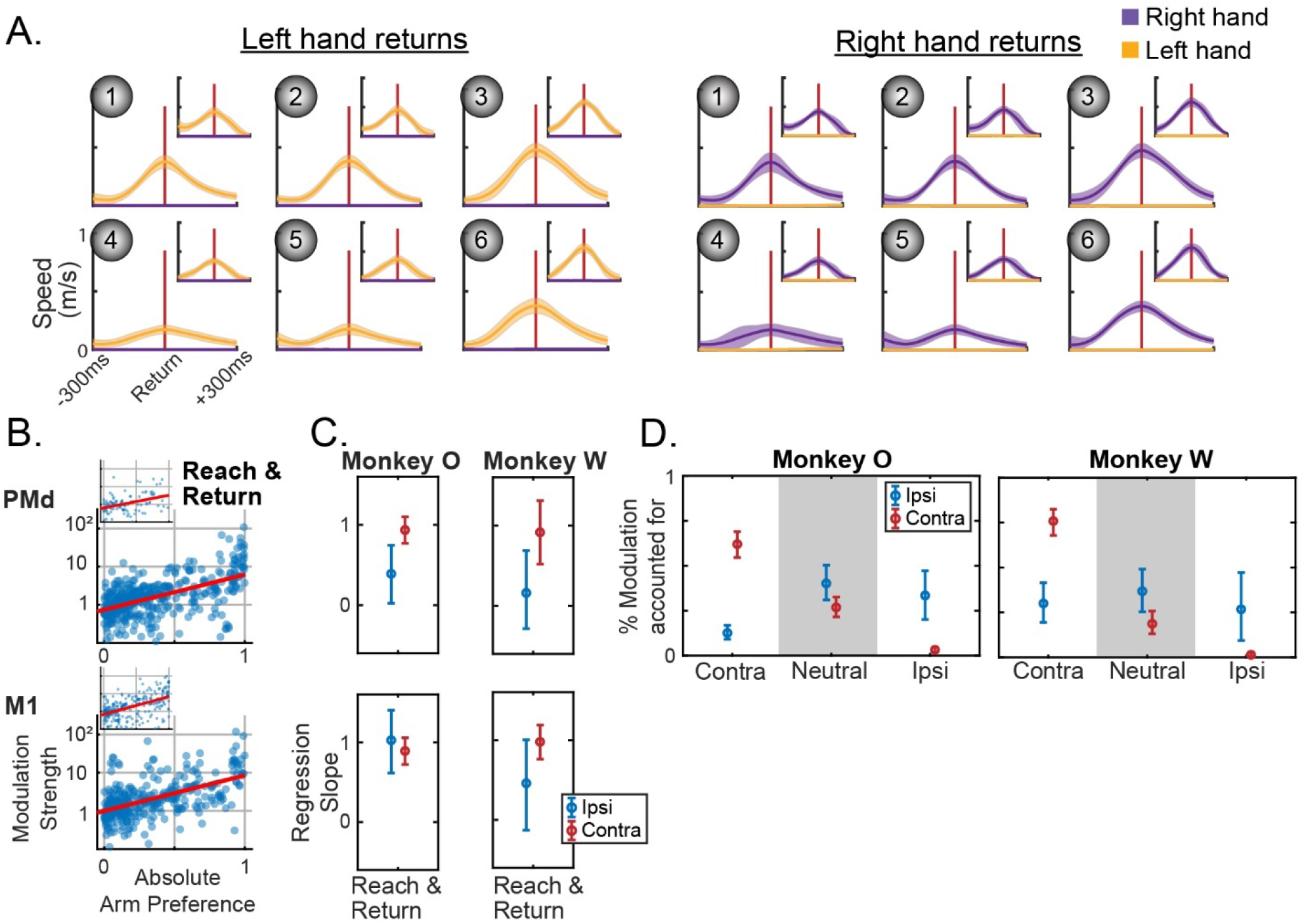
Dedicated signals persist with increased kinematic range. To determine whether having a limited range of reach directions was responsible for the observation of arm-dedicated signals, select analyses were performed again on data that included return movements. By including these movements, which were opposite the direction of the outward reaches used in the primary analyses, the range of sampled behavior was greatly increased. (A) Speed profiles for return movements following target acquisition during left-or right-hand trials. Individual trials were aligned to peak return speed, indicated by the vertical red line. Both reaching and stationary hands are plotted in each. Despite being unconstrained by the task, the non-selected hand remained still during the return. Monkey O main, monkey W inset. Mean +/− standard deviation. (B-D) Analyses from Figure 5 repeated using Move phase data concatenated with 300ms of data beginning 200ms before the point of peak return speed, i.e. reach and return. (B) Compare to Figure 5A. Modulation for the preferred arm plotted against arm preference, for all units in each brain area. Log-linear best fit lines are displayed in red. Inset figures belong to Monkey W. (C) Compare to Figure 5B. Slopes of regression lines fit to data from (B), independently for ipsi-and contra-preferring sub-populations. Mean +/− bootstrapped 95% confidence interval. (D) Compare to Figure 5F. The proportion of modulation within each partition from (C) during ipsi-or contralateral movements. Note that the total modulation is significantly lower for ipsilateral movements, particularly for Monkey W, and these data are only displayed as proportions. Mean +/− bootstrapped 95% confidence interval.

**Figure S4.**
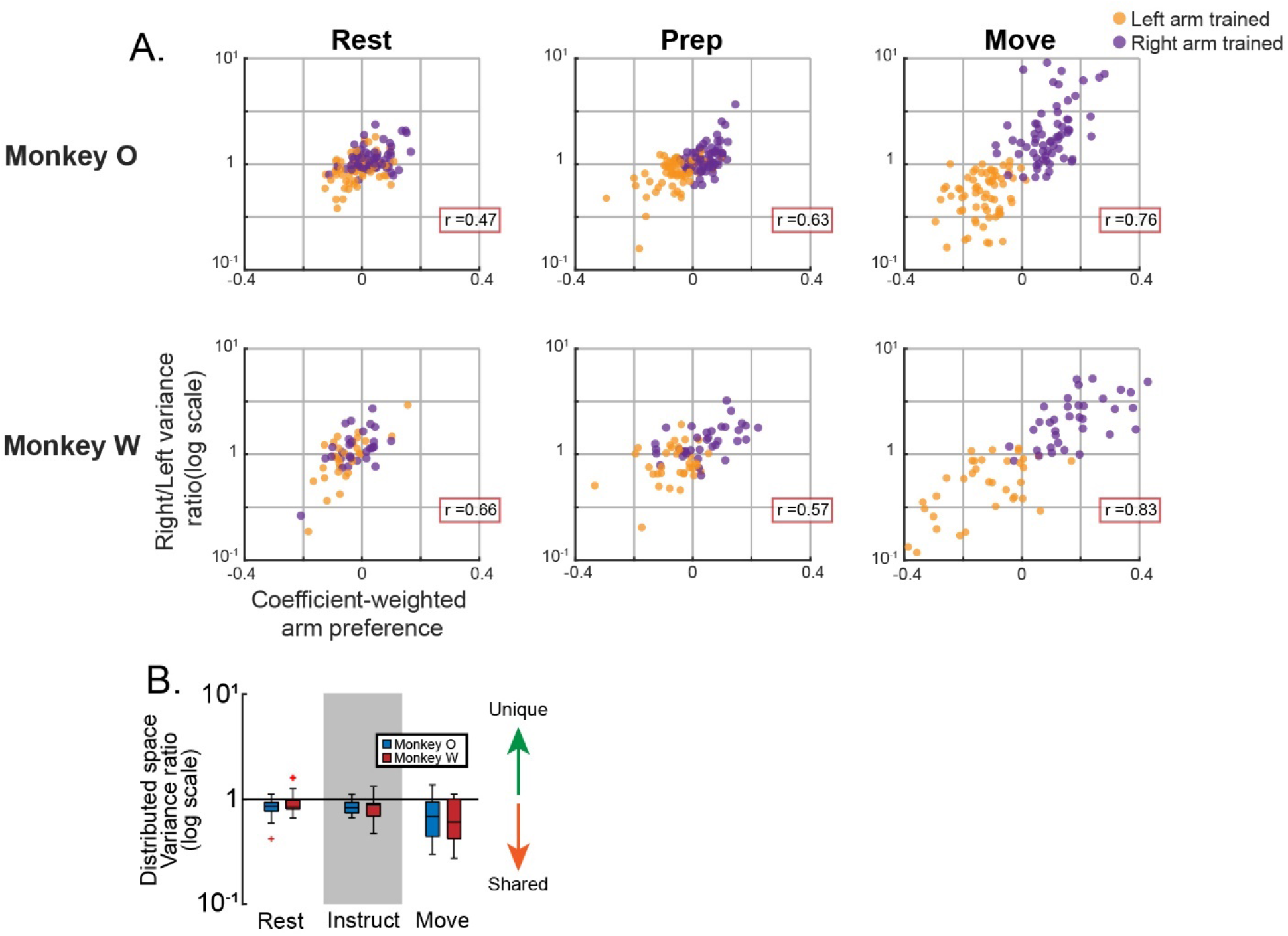
Subspace analysis using alternative firing rate normalization. Prior to performing PCA, an alternative method of normalizing firing rates was used for these plots. Rather than dividing by the standard deviation at Rest, each unit’s firing rate trace was divided by the full firing rate range + 5Hz (Elsayed et al., 2016; Ames and Churchland, 2019; Heming et al., 2019). This will mitigate the effect of highly modulated units, which PCA will preferentially represent otherwise. (A) Repetition of Figure 7D. (B) Repetition of Figure 8F.

**Figure S5.**
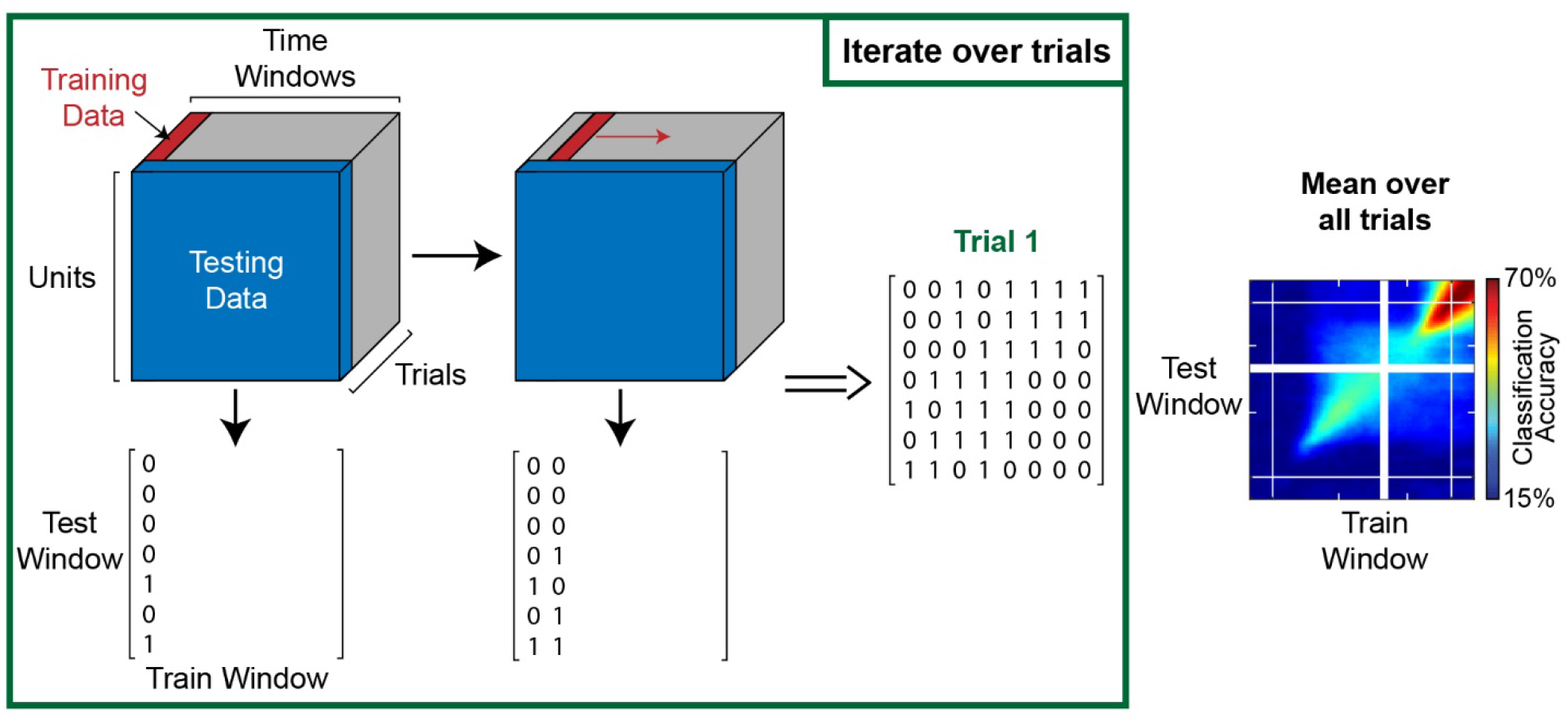
Method for fine timescale analysis of population coding and subspace separation. This schematic outlines the process for fine timescale analysis of population coding using LDA and leave-one-out cross-validation. Neural data were organized as 3D tensors (units, time windows, trials). Models were trained to predict targets using a single time window and all but one trial. Those models were then used to predict the target on the held-out trial, making separate predictions based on neural data from each time window. The process was then repeated using the next time window as training data until all possible pairs of time windows had been used as training and testing data. This constituted a 2D matrix of “hit” booleans (number time windows x number time windows) for the predictions of a single trial. After iterating over all trials to be used as held-out test data, the mean was taken across trials to construct a single 2D matrix of classification accuracy. The same basic process was used for visualizing the development of subspace separation, but instead of leave-one-out cross-validation trial sets were repeatedly divided into two random halves of equal size. Covariance alignment was then computed between all possible pairs of timepoints for the two disjoint trial sets.

## ACKNOWLEDGEMENTS

We thank M. Kitano for help in NHP care and handling, and A. You, E. Formento, W. Liberti and Z. Balewski for helpful discussions. This work was supported by the National Defense Science and Engineering Graduate Fellowship to T.C.D, and a grant from the National Institute of Health (NS097480) to J.M.C.

## AUTHOR CONTRIBUTIONS

T.C.D., C.M.M., R.B.I., and J.M.C. conceived and designed the experiments. T.C.D. performed the experiments, analyzed the data, and wrote the manuscript. T.C.D., C.M.M., J.D.W., R.B.I., and J.M.C reviewed and edited the manuscript.

